# A *Plasmodium falciparum* redox survival mechanism licenses killing by artemisinins

**DOI:** 10.1101/2025.10.08.681084

**Authors:** Smritikana Dutta, Mohammad Faaiz, Saikat Bhattacharjee, Kasturi Haldar, Souvik Bhattacharjee

## Abstract

Mutations in *Plasmodium falciparum* Kelch13 (K13) confer artemisinin resistance (ART-R) which threatens global malaria control, but known K13 functions fail to explain clinical ART-R. We reported that K13 binds the oxidant heme *in vitro*, however, its functions in redox-stress, cell survival and death remained unknown. Since taut control of free heme is not feasible in infected erythrocytes, we utilized a non-erythroid cell model to show that K13 directly binds and is stabilized by nanomolar heme levels. K13 also binds and regulates a major redox transcription factor, which is displaced by heme into the nucleus, to raise redox-stress responses that become suppressed during artemisinin-induced death (ART-death). K13’s evolutionarily conserved kelch domain confers heme-binding and ART-death characteristics to its mammalian orthologue KEAP1. Chemical or genetic elevation of K13, fuels ART-death proportionate to K13 levels even in vast excess of heme, suggesting a novel plasmodial redox-survival mechanism licenses ART-death in clinical ART-R.

## Introduction

Artemisinins (ARTs) are highly effective in curing blood-stage infection of *Plasmodium falciparum,* the deadliest of human malaria parasites^1^. These drugs are critical components of artemisinin-based combination therapies (ACT) that have played a key role in reducing the global burden of malaria and remain the cornerstone of its control worldwide^2^. Despite the prevalent model that ART kills through indiscriminate alkylation and damage of parasite lipids, proteins and DNA^3–5^, robust clinical resistance has been ascribed to one major gene *Pfkelch13* (K13; PlasmoDB ID: PF3D7_1343700)^6^, which has been shown to be causal for resistance^7^. Since ART-resistance (ART-R) poses a significant threat to malaria control^8,9^, K13 has been intensely studied over the last decade^10–14^. The consensus data suggest that it possess an evolutionarily conserved ‘kelch domain’ is cytoplasmically synthesized and associated with vesicular structures, including the parasite’s digestive endosomes, apicoplast, mitochondria, endoplasmic reticulum as well as vesicles enriched in Rabs, ATG18 and phosphatidylinositol 3-phosphate (PI3P)^9–11^. At the parasite’s trophozoite stage, when haemoglobin uptake and digestion are most active, K13 regulates endocytosis *via* structures called cytostomes, and resistance mutations suppress this process^15,16^. But notably, these very trophozoite stages harbouring K13 resistance mutations remain highly susceptible to ART suggesting that while hemoglobin uptake regulated by K13, it may not be the major target of resistance, which may arise from changes in additional but still unknown functions of K13.

ARTs are ‘pro-drugs’ in that they must all be converted to dihydroartemisinin (DHA) and which in turn is subsequently activated to release their killing action. High (micromolar) concentration of heme have been shown to increase ART-potency because it cleaves the endoperoxide bridge yielding highly reactive oxy-centered and C4-centered radicals^17^ that alkylates a wide range of cellular components causing oxidative damage, which results in parasite cell death^18,19^. Malaria parasites digestion of host hemoglobin actively produces heme in the parasite’s food vacuole (FV). Heme is presumed to confer the sensitivity of ARTs in killing malaria parasites^20,21^. However, it is poorly understood whether this can be achieved by physiological heme concentrations and how killing is effected in ‘ring’ parasites early in blood-stage infection that lack a food vacuole, yet are critical clinical targets of ART-R^22,23^.

We have recently reported that recombinant and parasite-derived K13 directly bind molecular heme with a dissociation constant (K_d_) of 61.5 nM^24^. Heme binds to many proteins^25,26^. This includes albumins with a K_d_ range of 2-40 µM^27^, explaining their ready capacity to transfer heme to wide range of acceptors^28^. In contrast heme-oxygenases which degrade heme, bind it with nanomolar (3-300 nM) affinity^29^. This suggests that with a K_d_ of 61.5 nM, K13 engages in at least moderately tight heme-binding. Heme binding is known to effect profound structural changes in proteins^30^ and heme is implicated in oxidative stress metabolism^31^. However, the consequences of heme binding for K13’s stability and turnover, and whether heme-binding induces K13-mediated redox-controlled responses remain entirely unknown. Furthermore, K13’s engagement of antioxidant and detoxification systems, as regulated by its closest mammalian orthologue Kelch-like ECH-associated protein 1 (KEAP1)^6,32,33^ has not been investigated. Finally, the implications of dislodging or neutralizing heme bound to K13, for mechanisms of ART-based killing remain to be discovered.

Since *in vitro* studies suggest that K13 binds heme with a K_d_ of 61.5 nM^24^, regulation of associated cellular mechanisms may be expected to be initiated at nanomolar to low micromolar concentrations. However, carefully controlling heme levels in this range remains unfeasible in blood-stage parasites. This is because the host erythrocyte contains high levels of hemoglobin (3.2-3.6 mM)^34^, whose digestion is a major source of parasite heme. Erythrocytes also contain high levels (21 ± 2 µM) of free heme^35^ which may translocate through passive and facilitated pathways^26,36^. Since mature human red cells are enucleated and terminally differentiated, these host sources of heme cannot be genetically ‘turned off’ post infection. Moreover, their reduction compromises parasite health^37^. Genetic or chemical targeting of parasite’s *de novo* heme biosynthesis in infected erythrocytes appears to have no effect on parasite growth^38,39^, suggesting that heme provided by exogenous pathways can be supplemented from host-derived sources. Notably, free heme levels in most non-erythroid cells are in the nanomolar range (25-300 nM)^36^, but in *P. falciparum* they are reported to be considerably higher (at 1.6 µM or more)^40,41^, suggesting that throughout blood-stage infection the parasite may reside in a high intracellular heme-rich environment, but the consequences for redox stress remain obscure.

We desired a cell system to study the molecular functions of K13 and their regulations by nanomolar concentrations of heme. We therefore used a model non-erythroid mammalian cell line Du145^42^ with reduced levels of KEAP1, adapted to grow in heme-depleted media, and whose free heme levels were tightly regulable by blocking endogenous heme biosynthetic enzymes. These Du145 cells were otherwise capable of an active redox response^43^. Our goal was to query whether K13 and/or heme contribute to redox-stress responses well as obtain mechanistic understanding of how either or both trigger sensitization of cells to killing by ARTs and validate the model system as a rapid, functional screen to identify redox cell survival and death mechanisms of *Plasmodium falciparum* malaria

## Results

### K13 cellular levels are increased by heme-binding and -stabilization

Human Du145 cells (HTB-81, ATCC, www.atcc.org) contain a full length wild-type *Keap1* gene, but CpG dinucleotide methylation of its core promoter results in low *keap1* mRNA expression^44^, making them suitable for ectopically expressing kelch domain proteins with minimal interference from the native KEAP1 protein. Sequence comparison clearly revealed that K13 is a homologue of KEAP1 (**Supplementary Fig. 1a**). We generated three stable transgenic lines expressing: (i) K13 as a C-terminus fusion to GFP; K13-GFP, (ii) GFP alone, as well as (iii) a C-terminus fusion of flag-strep tag KEAP1 (see *Materials and Methods*). Indirect immunofluorescence assays (IFA) with quantitative analyses, suggested cellular distribution of K13-GFP, GFP and KEAP1, at comparable levels in transgenic Du145 cells (**Supplementary Fig. 1b)**. K13-GFP, GFP and flag-strep tag KEAP1 respectively appeared as fusion proteins at expected sizes of 105-kDa, 27-kDa and 80-kDa (**Supplementary Fig. 1c, d**).

We recently showed that *in vitro* heme bound recombinant K13 protein (TrK13-WT) with high affinity^24^, but action of heme in cells was not known. Since the K_d_ for heme binding to K13 protein was 61.5 nM, we wanted to measure the effects at low nanomolar concentrations of heme. This required growing Du145 cells under conditions where available heme was substantially depleted from the extracellular sources, and the production of biosynthetic heme was blocked by inhibitors. Fetal bovine serum (FBS) is the largest extracellular donor of heme and contributed median concentration of 3.2 ± 0.06 µM heme to culture media. We reduced FBS concentrations of heme to <850 nM (**Supplementary Fig. 2a, b**) to create heme-depleted media (HDM; *Material and Methods*). Since the K_d_ of heme for albumins is >1 µM, this was expected to dramatically reduce the transfer of extracellular heme to cells. We also added succinylacetone [SA; an inhibitor of the second enzyme of heme biosynthesis pathway, δ-aminolevulinic acid (ALA) dehydratase] to the culture media, which restricted intracellular *de novo* heme synthesis. Our overall goal was to achieve culture conditions that would be responsive to nanomolar concentrations of exogenously added heme. Accordingly, under HDM conditions, the parental Du145, transgenic GFP and K13-GFP cells all showed intracellular concentrations of ∼60 nM heme. Addition of extracellular heme at 5 µM, raised intracellular heme to 400-450 nM. When extracellular concentrations were raised to 50 µM, the intracellular levels plateaued at ∼750 nM (**Supplementary Fig. 2c**). Together these data suggested that in the course of all experiments intracellular heme concentrations ranged from ∼60-750 nM.

K13-GFP cells cultured in HDM (K13-GFP^HDM^) were subsequently exposed to different concentrations of exogenously added heme (0-5 µM) in the media (referred to as heme-replenished media). As shown in **Fig. 1a, concentrations** of 10 nM heme induced detectable increase in K13-GFP levels. Progressive increase of K13-GFP levels were seen at 50 nM, 100 nM, 1 µM and 5 µM heme. At 5 µM heme, K13-GFP levels were restored to those seen in K13-GFP ‘control’ cultures (**Fig. 1b**). There was no effect on the cellular viability of K13-GFP (or GFP) expressing cells at 5 µM heme; (**Supplementary Fig. 2d**: the IC_50_ for heme was ∼25-fold higher for all cells). Notably, there was no detectable change in the relative intensity of GFP or KEAP1 proteins when cells were subjected to heme depletion or when supplemented with 5 µM heme (**Supplementary Fig. 2e, f**).

**Fig. 1.**
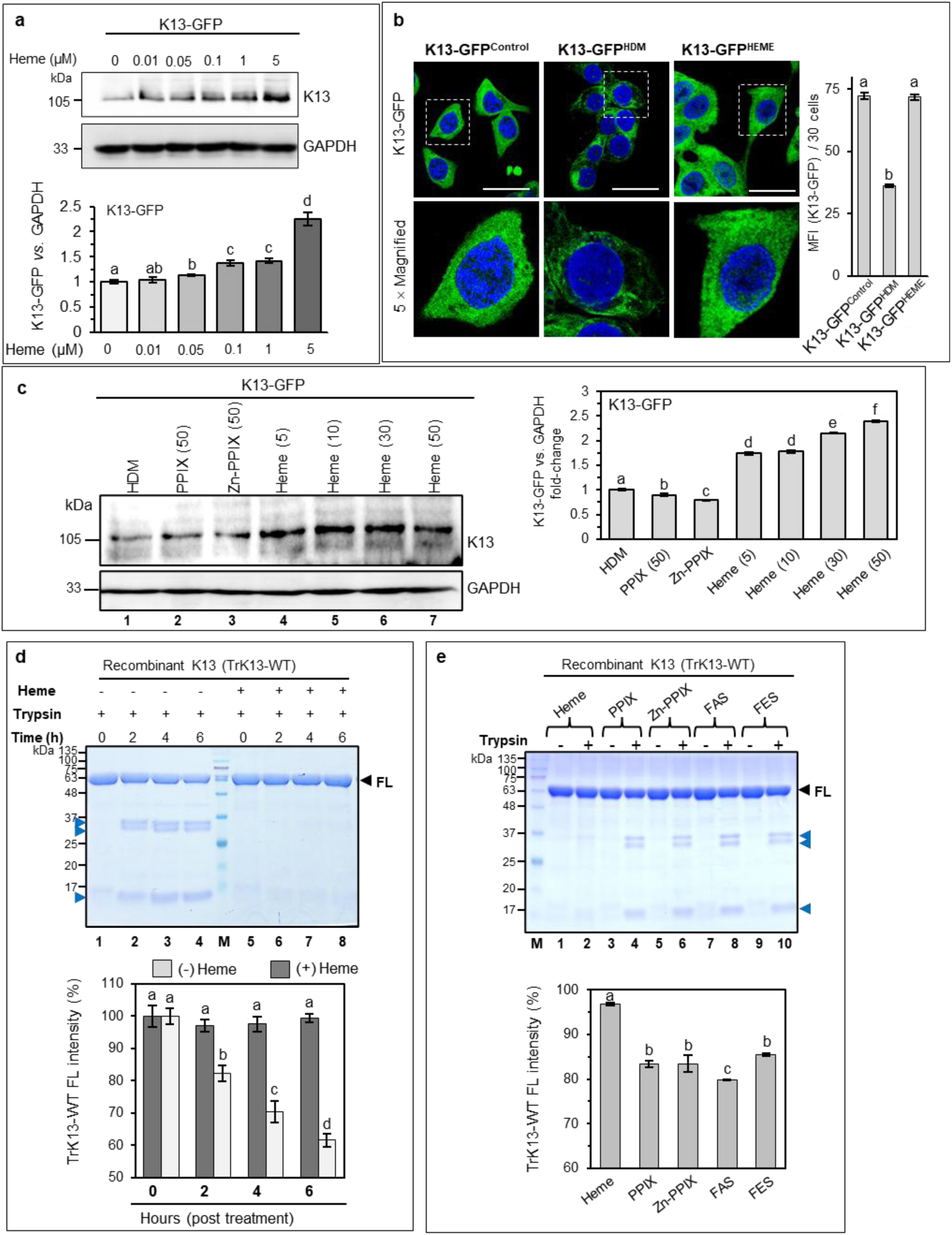
Heme stabilizes K13-GFP and confers *in vitro* resistance to trypsin digestion. **a.** Western blots (top) and densitometric analysis (bottom) for K13-GFP under HDM condition or when supplemented with increasing concentrations of heme. GAPDH is the loading control. **b.** IFA images (left) and mean fluorescence intensity plot (MFI, right) of K13-GFP cells under control, HDM or 5 µM heme-supplemented conditions. Scale bar, 5 µm. **c.** Western blot (left) and quantitative densitometric analysis (right) of K13-GFP band intensity under HDM condition (lane 1), or upon supplementation with heme analogs PPIX (lane 2), Zn-PPIX (lane 3) or increasing concentrations (µM) of heme (lanes 4-7). Values in parenthesis indicate concentration (in µM). **d.** Coomassie-stained SDS-PAGE showing trypsin-digested fragments (blue arrowheads) derived from the full length (FL, black arrowhead) recombinant K13 (TrK13-WT) in heme-free (lanes 2-4) and heme-bound TrK13-WT (lanes 6-8) protein. The heme-free and heme-bound TrK13-WT proteins without trypsin treatment are shown in lanes 1 and 5, respectively. **e.** Coomassie-stained SDS-PAGE showing trypsin-digested fragments (blue arrowheads) derived from the recombinant FL TrK13-WT protein (black arrowhead) when preincubated with heme analogs PPIX, Zn-PPIX, FAS or FES (lanes 4, 6, 8 and 10, respectively) and not by the heme-bound TrK13-WT (lane 2) protein. Lanes 1, 3, 5, 7 and 9 represent samples with no trypsin treatment. Molecular weight standards (M) for all SDS-PAGE and western blots (in kDa) are as indicated. Graphs represent mean from three biological replicates ± SE. Statistical significance calculated by one-way Anova. Distinct alphabets represent significant difference at p.adj ≤ 0.05.

We next investigated the specificity of heme action. As shown in **Fig. 1c, PPIX** alone or when complexed with Zn, had no effect even at 50 µM. However, increasing heme levels from 5 to 50 µM continued to steadily increase levels of K13-GFP. Together, the data suggest that heme at extracellular concentrations of 10 nanomolar or higher (and up to 50 µM) increase cellular levels of K13-GFP (but not GFP or KEAP1), suggesting it is specific for K13. Increases in K13 were only mediated by heme (not PPIX or Zn-PPIX) and likely due to heme’s directly binding to K13 (since we have previously shown that heme binds K13 with nanomolar affinity *in vitro*^24^).

Amongst heme binding proteins, hemopexin, a serum glycoprotein (which displays highest affinity for heme at K_d_ ∼ 1 pM^45,46^) has been reported to show heme-dependent resistance to protease digestion^45^. We therefore investigated if the protease sensitivity of K13 was altered in its heme-bound state. For *in vitro* assay, we used the affinity purified 61-kDa recombinant TrK13-WT protein as K13 representative, described in our previous studies^24,47^. Recombinant TrK13-WT protein (10 µM) was incubated with or without equimolar concentrations of heme and desalted to remove excess unbound heme (see *Materials and methods*). Heme-bound TrK13-WT protein was subsequently treated with trypsin under optimal buffer and pH conditions for varied incubation times. For trypsin (a serine protease that cleaves between the carboxyl group of arginine or lysine and the amino group of the adjacent amino acid), we also included commercially procured BSA, a confirmed heme-binding protein^48^ (but with substantially lower affinity) as a control. As shown in **Fig. 1d, Coomassie**-stained SDS-PAGE revealed full-length (FL) TrK13-WT protein at 61-kDa. Treatment with trypsin yielded additional low molecular weight fragments (doublets at 30-kDa and at 12-kDa; blue arrowheads) in the absence of heme, whose appearance was blocked when the TrK13-WT protein was pre-incubated with heme. Heme-binding only conferred partial resistance of BSA to trypsin (**Supplementary Fig. 2g**). In absence of heme, multiple small protein fragments were detected (from 22-40 kDa; green arrowheads), while in its presence, larger fragments were observed at 45-55-kDa (red arrowheads). The effects of heme on trypsin digestion of both TrK13-WT protein and BSA were likely because of changes in the 3-dimensional conformation of these proteins (which is consistent with previous reports in the literature^48^ and observed in our earlier CD studies^24^). As shown in **Fig. 1e, resistance** to trypsin digestion was not seen when the TrK13-WT protein was incubated with PPIX or Zn-PPIX or iron salts (ferrous ammonium sulfate, FAS or ferric ammonium sulfate, FES, although these salts are known to bind K13)^24^. Together, these data suggest that heme binding to K13 shows chemical and molecular specificity, may induce confirmational change and thereby induce stabilization of K13 in cells.

### K13-regulated oxidative stress responses are triggered by heme

Nuclear factor erythroid 2-related factor 2 (Nrf2) is a major redox-regulated transcription factor controlled by KEAP1^49^, the closest mammalian orthologue of K13. We therefore assessed whether Nrf2 was regulated by K13 and the impact of heme, a known, major redox regulator^50^. First, we investigated changes in Nrf2 levels in GFP or KEAP1 expressing cells under HDM or heme-replenished culture conditions *i.e.,* GFP^HDM^ *versus* GFP^HEME^ or KEAP1^HDM^ *versus* KEAP1^HEME^ cells. Relative intensities of both GFP and KEAP1 proteins remained unaffected by heme (**Supplementary Fig. 3a, b**). Consequently, we investigated only K13-GFP cells, cultured in the absence or presence of increasing concentrations of heme, and followed by immunoprecipitations from cellular lysates using anti-GFP antibodies. Western blot (**Fig. 2a**) and quantitative densitometric analysis (**Fig. 2b**) of band intensities revealed increasing amounts of immunoprecipitated K13-GFP with increased heme concentrations (**2a i** and **2b i**). In contrast, levels of co-immunopurified Nrf2 decreased with increasing heme (**Fig. 2a i**), although there was increase in the overall cellular Nrf2 levels (**Fig. 2a ii** and **2b ii-iii**). These results strongly implied that interactions between K13 and Nrf2 were negatively affected by heme. To investigate whether these reflected direct protein-protein interactions, we co-incubated recombinant Nrf2-6×his protein with either heme-free or heme-bound recombinant TrK13-WT (that was desalted to remove excess unbound heme). As shown in **Fig. 2c, after** incubation with heme for 1 h, there was ∼26% reduction in Nrf2-6×his binding to TrK13-WT, suggesting heme may directly prevent/disrupt Nrf2 binding to K13.

**Fig. 2.**
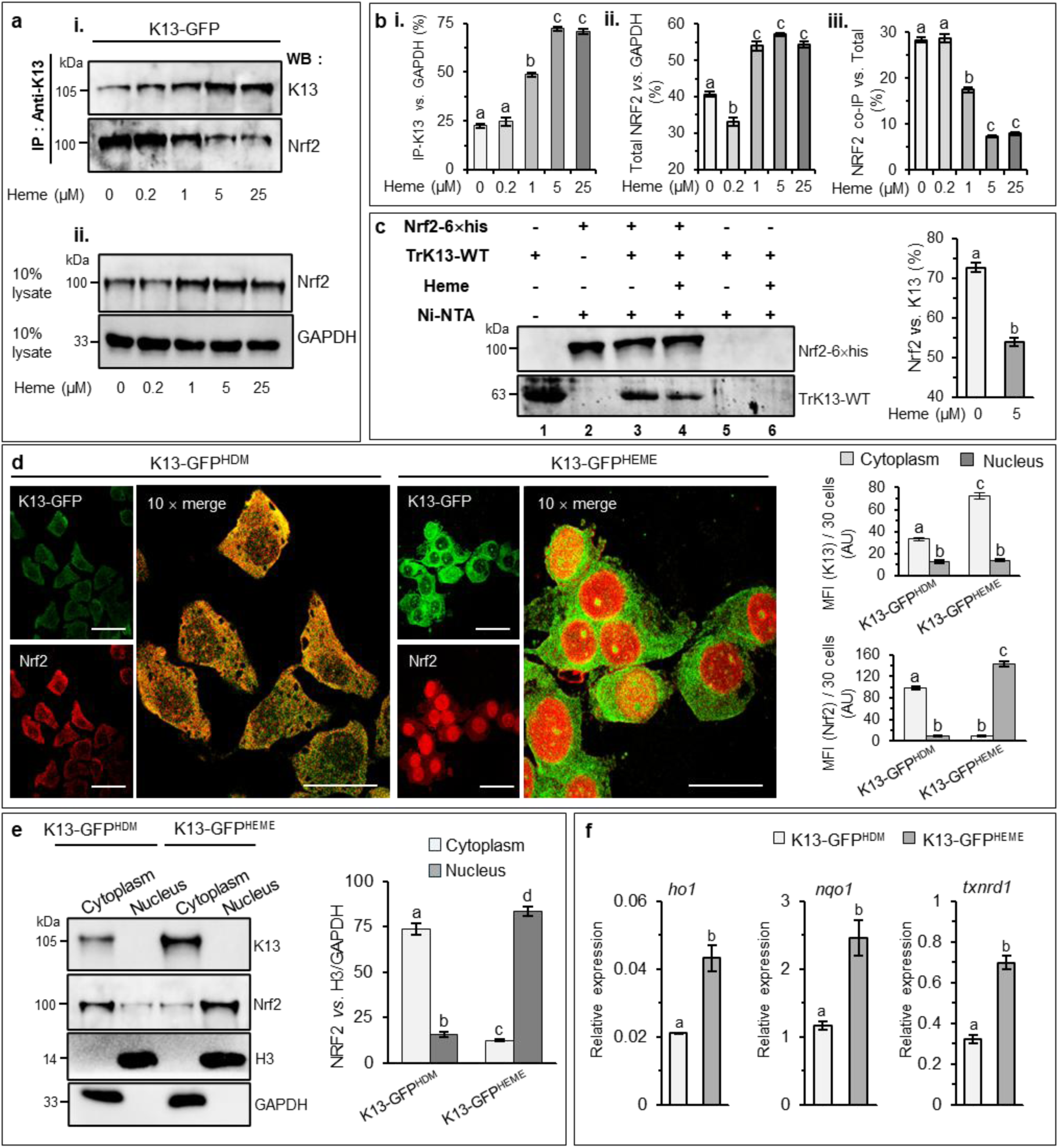
Heme alters the level and dynamics of the redox reporter Nrf2 in K13-GFP-expressing cells. **a. i.** Western blots showing immunoprecipitated K13-GFP (top) and the co-associated Nrf2 (bottom) protein from K13-GFP cells cultured in HDM or in the presence of different concentrations of heme. **ii.** Total Nrf2 (top) and GAPDH (bottom, as loading control) in the cellular lysates for conditions in a. **b.** Graphical densitometric analyses of K13-GFP, total Nrf2 and co-purified Nrf2 for data shown in a. **c.** Western blots (left) and graphical densitometric analysis (right) showing reduced co-purification of the heme-bound (lane 4) recombinant K13 (TrK13-WT) by the recombinant Nrf2-6×his protein (pre-bound to Ni-NTA beads) as compared to the heme-free TrK13-WT protein (lane 3). Lanes 1 and 2 indicate TrK13-WT input and the Ni-NTA bound Nrf2-6×his protein, respectively. Lanes 5 and 6 show no nonspecific binding of heme-free or heme-bound TrK13-WT protein to the Ni-NTA beads, respectively. **d.** IFA images showing increased levels of K13-GFP (green) fluorescence and nuclear localization of Nrf2 (red; right panel) in the K13-GFP^HEME^ cells (right). In K13-GFP^HDM^ cells (right), the Nrf2 fluorescence distribution was predominantly cytoplasmic. Scale bar, 5 µm. **e.** Western blots and graphical densitometric quantitation for cytoplasmic and nuclear fractionation of Nrf2 in K13-GFP^HDM^ and K13-GFP^HEME^ cells. **f.** RT-qPCR data showing relative expression of stress-response genes in K13-GFP^HDM^ cells and K13-GFP^HEME^ cells. Molecular weights (in kDa) are as indicated. Error bars represent mean of three technical replicates ± SE. Statistical significance was calculated by one-way Anova. Distinct alphabets represent significant difference at p.adj ≤ 0.05.

We next performed immunolocalization studies to determine subcellular distribution of Nrf2 in K13-GFP^HDM^ and K13-GFP^HEME^ cells (**Fig. 2d**). Here, data revealed predominantly cytoplasmic localization of Nrf2 with K13-GFP under HDM conditions (left panel). In contrast, the K13-GFP^HEME^ cells displayed prominent nuclear localized Nrf2 protein (right panel). Moreover, in the presence of heme, levels of K13 were higher than under HDM conditions. Subcellular fractionation confirmed that under conditions of HDM, a vast majority (∼83%) of Nrf2 protein was detected in the cytoplasmic fraction (**Fig. 2e**; second blot). However, in the K13-GFP^HEME^ cells, ∼87% of the Nrf2 protein was detected in the nuclear fraction. As expected, the K13-GFP protein was present exclusively in the cytoplasmic fraction independent of the absence or presence of heme. Furthermore, the intensity of K13-GFP was much higher under heme-replenished condition (top blot). The GFP expressing cells revealed no changes in the relative nuclear/cytoplasmic fractionation of the Nrf2 protein or intensities of GFP (**Supplementary Fig. 3c**). As Nrf2 is a redox-controlled transcription factor, we also compared the expression of stress-response genes *heme oxygenase* (*ho-1*), *NAD(P)H quinone oxidoreductase 1* (*nqo1*) and *thioredoxin reductase 1* (*txnrd1*) in K13-GFP^HDM^ and K13-GFP^HEME^ cells. As shown in **Fig. 2f, RT**-qPCR data confirmed increased expression of these stress-response genes as a consequence of increased Nrf2 translocation to the cell nucleus.

### Chaperone mediate autophagy is a major pathway of K13 degradation

A recent study implicated chaperone-mediated autophagy (CMA) in the lysosomal degradation KEAP1 in mouse midbrain progenitor cell line SN4741^51^. Since K13 shares structural and sequence identity with KEAP1, we investigated if similar degradation pathway was involved in the proteolysis of K13-GFP under HDM conditions. Our analyses of the primary sequence of K13, suggested it contained ‘KFERQ-like’ motifs as recognized by an online algorithm KFERQ finder v0.8 (https://rshine.einsteinmed.edu)^52^. These motifs are targeting signals of CMA. In CMA the chaperone heat shock 70-kDa (cognate) protein 8 (also known as HSPA8) binds ‘substrate’ proteins bearing ‘KFERQ-like’ motifs^53,54^. LAMP2A acts as a receptor on the lysosomal membrane. HSPA8 binds the cytosolic tail of LAMP2A^55^. This in turn unfolds the substrate and enables its translocation into lysosomes where it is subsequently degraded^56^. As shown in **Fig. 3a**, K13 contains eighteen ‘KFERQ-like’ motif, strongly suggesting it may be targeted to lysosomes. Immunolocalization studies using indirect IFA suggested moderate association between K13 and LAMP2A with Pearson’s correlation coefficient (PCC) of 0.59. Notably, heme raised the PCC to 0.79, suggesting it induced a high level of co-association of K13-GFP with LAMP2A (**Fig. 3b**). We therefore concluded that although heme induces conformational change in K13, this does not obscure targeting of K13-GFP to LAMP2A labeled lysosomes. But since heme increases levels of K13, it is not likely to promote degradation of K13-GFP within the lysosome. We therefore propose (as schematic in **Fig. 3b**) that the heme-induced conformational stability may prevent unfolding of the K13-GFP substrate and its translocation into the lysosome, thereby limiting its degradation in cells.

**Fig. 3.**
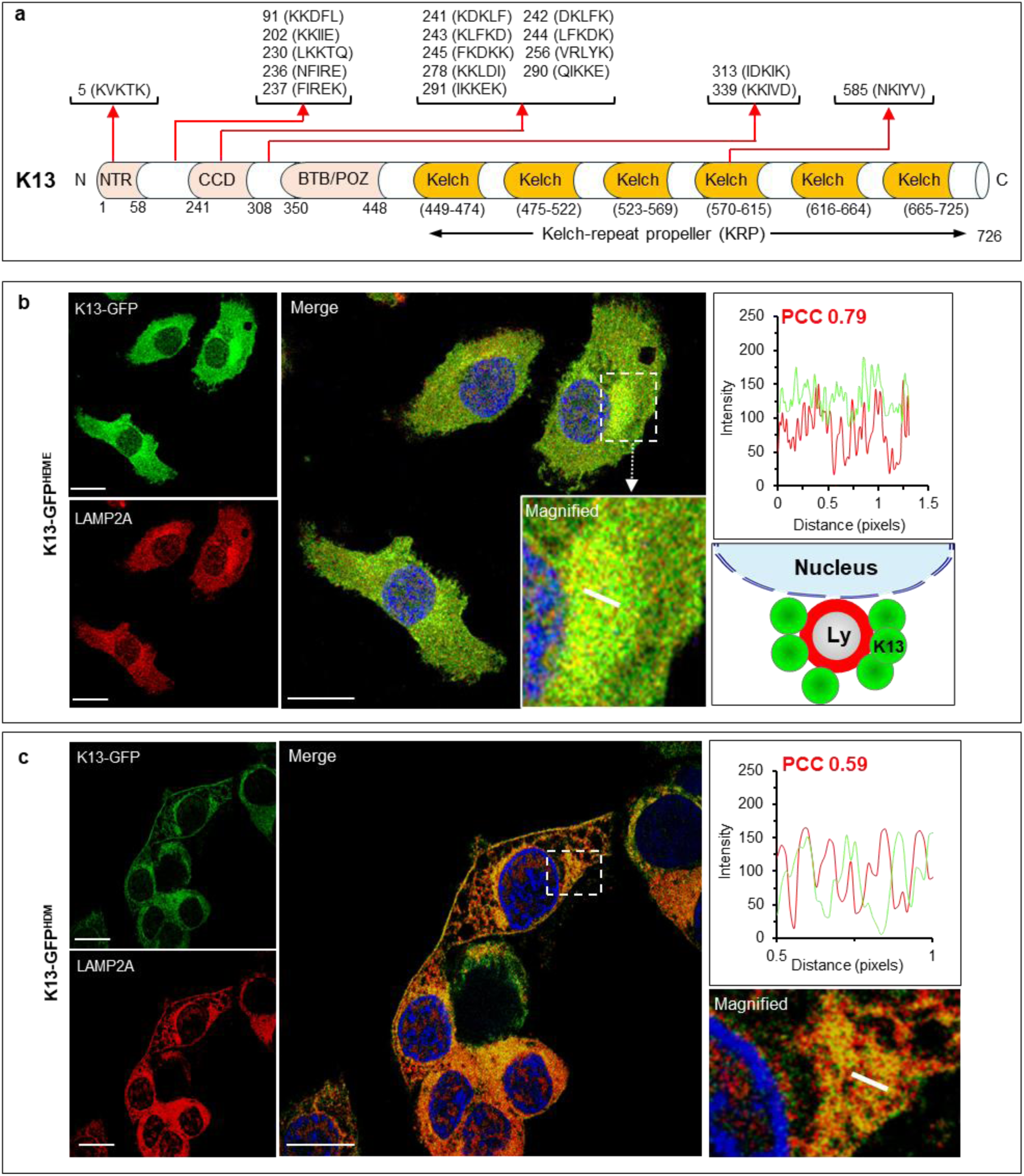
Signature motifs of autophagy in K13 and heme-dependent co-association with LAMP2A. **a.** Schematic showing the distribution of 18 putative KFERQ-like motifs in different domains of K13, as identified by KFERQ finder v0.8 (https://rshine.einsteinmed.edu). NTR, N-terminus domain; CCD, coiled-coil domain; BTB-POZ, Broad-complex, tramtrack and bric-a-brac (BTB)-poxvirus and zinc-finger (POZ) domain. **b-c.** IFA images (left) and RGB intensity plots (right) showing the extent of colocalization between K13-GFP (green) and LAMP2A (red) in K13-GFP^HDM^ (b) and K13-GFP^HEME^ cells (c). Pearson’s correlation coefficient (PCC) values for the selected region (white bar) are indicated in red text. Magnified images of the white-dotted boxed region for respective cells are also shown. Schematic in b represents proposed association of K13 with lysosomes based on the inset magnification. Scale bar, 5 µm is shown at the bottom of each image. Cell nucleus (blue) stained with Hoechst 33342.

Since LAMP2A is an exclusive CMA marker, we investigated the effects of a previously reported CMA-inhibitor, *i.e*., the RARα (retinoic acid receptor alpha) activator ATRA (*all-trans* retinoic acid), known to be a potent inhibitor of LAMP2A expression^57^. As expected, under HDM conditions, there was moderate co-association of K13 and LAMP2A, with a PCC of 0.58 (**Fig. 4a i**). Treatment with ATRA elevated levels of K13-GFP as well as its co-association with LAMP2A (raising the PCC to 0.90; **Fig. 4a ii**). As expected, heme also increased K13-GFP levels and the PCC with LAMP2A rose to 0.77 (**Fig. 4a iii**). Notably, ATRA markedly increased K13-GFP in HDM to levels far higher than those seen with heme replenishment (**Fig. 4b i** and **4b iii**). ATRA also significantly reduced LAMP2A levels compared to controls, both in K13-GFP^HDM^ and K13-GFP^HEME^ cells (**Fig. 4b i** and **4b ii**). Together these data strongly suggested ATRA-mediated reduction of LAMP2A prevented the degradation of K13, explaining why it increased K13 levels more than heme.

**Fig. 4.**
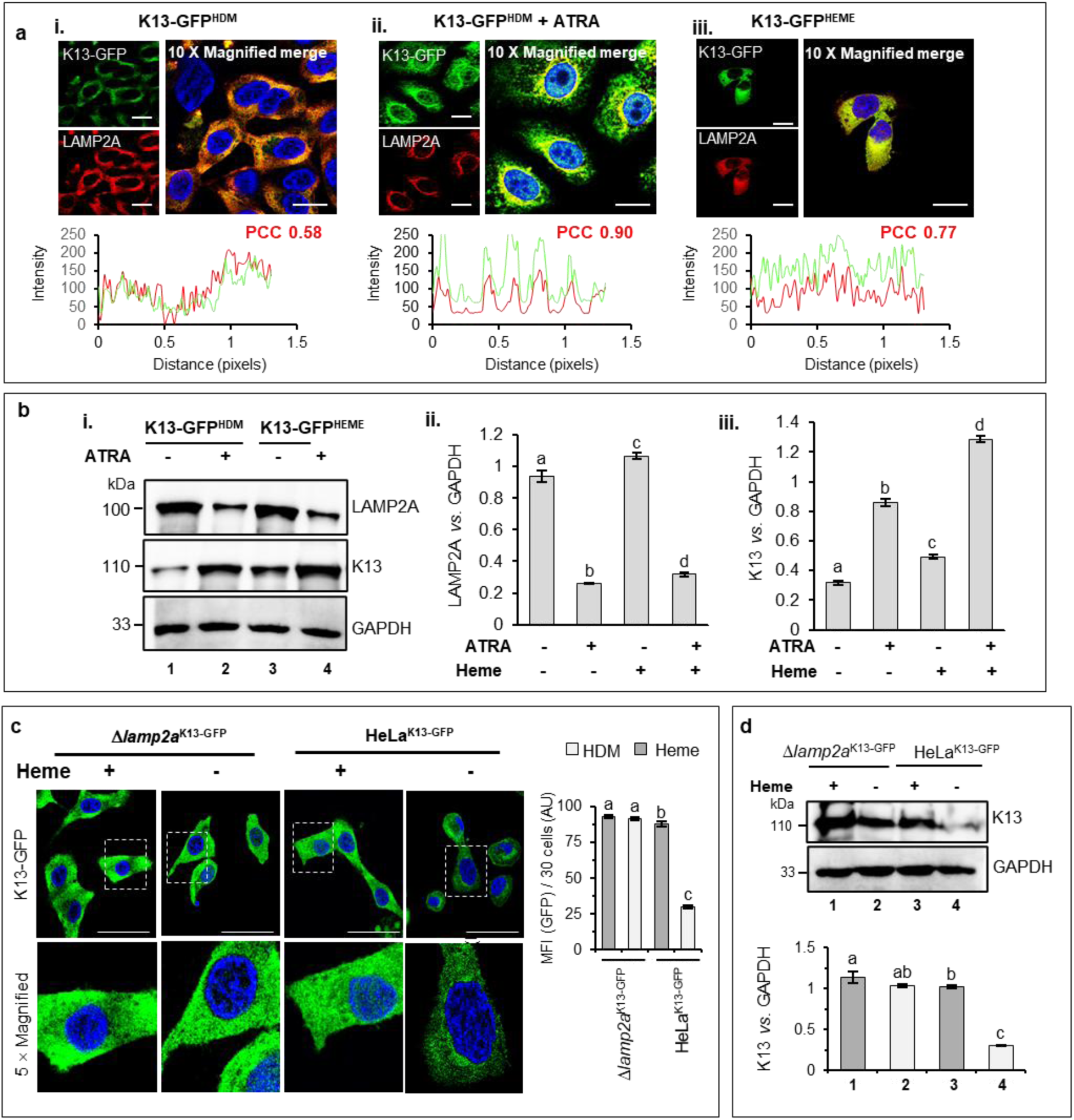
Reduction or knockout of LAMP2A decreases degradation of K13-GFP under HDM conditions. **a.** In Du145 cells, IFA images and RGB intensity plots (with PCC values) showing relative colocalization between K13-GFP (green) and LAMP2A (red) in K13-GFP^HDM^ cells in the absence (i) or presence (ii) of ATRA (inhibitor of LAMP2A expression) compared to K13-GFP^HEME^ cells (iii). Corresponding fluorescence and merged images were acquired under identical optics and exposure conditions. **b.** Western blots (i) and fold change (ii-iii) for LAMP2A (top), K13-GFP (middle) in the absence or presence of ATRA in cells expressing K13-GFP^HDM^ (lanes 1-2) or K13-GFP^HEME^ (lanes 3-4). GAPDH (bottom) was the loading control. **c.** In HeLa cells, IFA images (left), and graphical MFI (right) of K13-GFP fluorescence (green) in transgenic *Δlamp2a*^K13-GFP^ and parental HeLa^K13-GFP^ cells under HDM (−) or heme-supplemented (+) conditions. **d.** Western blots (top) and graphical fold change of K13-GFP (top) in transgenic *Δlamp2a*^K13-GFP^ (lanes 1-2) and parental HeLa (lanes 3-4) cells under HDM (lanes 2 and 4) or heme-supplemented (lanes 1 and 3) conditions. GAPDH was the loading control. Scale bar for IFA images, 5 µm. Boxed (white dotted) images are magnified in sub-panel. For all statistical analyses, each bar denotes the mean from three technical replicates ± SE. Statistical significance was calculated by one-way Anova. Distinct alphabets represent difference at p.adj ≤ 0.05. Molecular weights (in kDa) for all blots are as indicated.

To further confirm the role of LAMP2A-mediated degradation of K13-GFP, we obtained stable transgenic cell lines expressing K13-GFP or GFP in the HeLa *lamp2a* knockout (*Δlamp2a*; Abcam; ab255402) and the corresponding parental HeLa cells (Abcam, ab255928). HeLa cells have previously been reported to inherently express very low levels of native KEAP1 protein and correspondingly have high basal Nrf2 activity^58^. We designated them as *Δlamp2a*^K13-GFP^, HeLa^K13-GFP^, *Δlamp2a*^GFP^ and HeLa^GFP^, cells, respectively. As shown in **Supplementary Fig. 4a**, LAMP2A protein was undetectable in *Δlamp2a* cells both by fluorescence microscopy and in western blots. Moreover, there was no difference in levels of K13-GFP in the transgenic *Δlamp2a*^K13-GFP^ cells regardless of whether they were cultured under HDM or 5 µM heme-supplemented conditions (**Fig. 4c**). In contrast, significant reduction in the K13-GFP level was evident in the parental HeLa^K13-GFP^ cells in HDM (compare lanes 3 and lane 4). In *Δlamp2a*^GFP^ cells, we observed no changes in the level of GFP (**Supplementary Fig. 4b**) when compared to the HeLa^GFP^ cells irrespective of HDM or heme-replenished conditions. Thus, the absence of LAMP2A adversely affected the proteolysis of K13-GFP, but not GFP, which further substantiated the involvement of CMA pathway in proteolysis of K13-GFP under HDM conditions.

### K13’s kelch domain transfers heme-dependent stability and heme binding property to KEAP1

We next investigated if the heme-regulated stability of K13 was due to its kelch repeat (KRP) domain and could be transferred to KEAP1. As shown in **Fig. 5a**, we made chimeras of the N-terminus of KEAP1 and the kelch domain of K13 fused to GFP (KEAP1^N^K13^KRP^-GFP) as well N-terminus of K13 and the kelch domain of KEAP1 fused to GFP (K13^N^KEAP1^KRP^-GFP). Stable expression of KEAP1^N^K13^KRP^-GFP and K13^N^KEAP1^KRP^-GFP proteins in Du145 cells were confirmed by western blots (**Fig. 5b**). As shown in **Fig. 5c**, immunolabeling assays revealed significant (72%) reduction in KEAP1^N^K13^KRP^-GFP cells under HDM conditions when compared to respective counterparts in 5 µM heme-supplemented media. In contrast K13^N^KEAP1^KRP^-GFP cells were reduced by only 30%. Supplementing HDM with the heme precursor ALA had no significant effect, but heme largely restored levels of KEAP1^N^K13^KRP^-GFP expression (**Fig. 5d)**. K13^N^KEAP1^KRP^-GFP showed much less heme-responsive expression (**Fig. 5e**). Together, these results suggested that the kelch domain of K13 conferred robust heme-stimulated expression of the KEAP1-K13 chimera which was closely comparable to the wild-type K13-GFP. The NTR, CCD and BTB-POZ domains of K13 also made a minor contribution in transferring heme-dependent phenotype to the KEAP1 chimera.

**Fig. 5.**
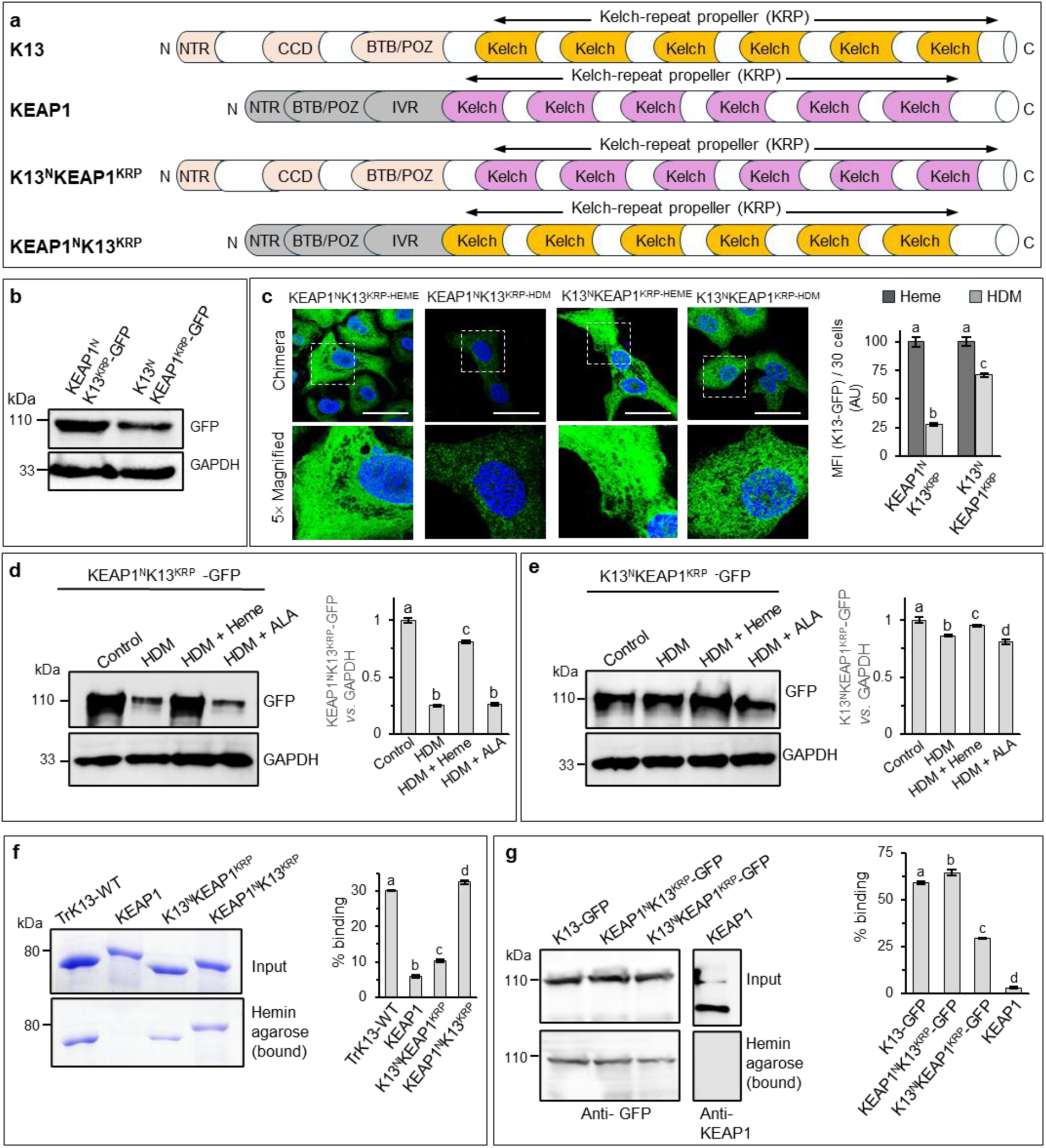
The K13 KRP domain confers heme-dependent stability and heme-binding characteristics to KEAP1. **a.** Schematics for domain organization in K13, KEAP1, K13^N^KEAP1^KRP^ and KEAP1^N^K13^KRP^ chimeric proteins. NTR, N-terminus domain; CCD, coiled-coil domain; BTB-POZ, Broad-complex, tramtrack and bric-a-brac (BTB)-poxvirus and zinc-finger (POZ) domain; IVR, intervening region. **b.** Western blot detecting K13^N^KEAP1^KRP^-GFP and KEAP1^N^K13^KRP^-GFP ^(^arrowhead) chimeras in transgenic Du145 cells. Loading control is GAPDH. **c.** IFA images and mean fluorescence intensity (MFI) of K13^N^KEAP1^KRP^-GFP and KEAP1^N^K13^KRP^-GFP cells cultured under HDM or heme-repleted conditions. Nuclei stained with Hoechst 33342 (blue), scale bar, 5 µm. Respective boxed (white dotted) regions are magnified and shown below. **d-e.** Western blots (left) and densitometric fold change (right) in KEAP1^N^K13^KRP^-GFP (d) and K13^N^KEAP1^KRP^ (e) protein band intensities under HDM, heme-repleted or heme precursor aminolevulinic acid (ALA)-containing HDM conditions. **f-g.** Coomassie-stained SDS-PAGE (f) and western blots (g) showing hemin agarose binding by indicated KEAP1-K13 recombinant proteins or chimeras (f) or from respective transgenic Du145 cellular lysates (g). Percentage binding (as compared to the total input) is shown graphically. Molecular weights (in kDa) for SDS-PAGE and western blots are as indicated. Error bars in the graphs represent mean of three technical replicates ± SE. Statistical significance was calculated by one-way Anova. Distinct alphabets represent significant difference at p.adj ≤ 0.05.

In addition, we investigated the relative affinity of recombinant K13 (also known as TrK13-WT), KEAP1, KEAP1^N^K13^KRP^ and K13^N^KEAP1^KRP^ proteins for hemin-agarose. Recombinant proteins expressed and purified from *E. coli* were incubated with hemin-agarose beads and percentage binding was determined (see *Materials and Methods*). As shown in **Fig. 5f**, K13/TrK13-WT, KEAP1, KEAP1^N^K13^KRP^ K13^N^KEAP1^KRP^ proteins respectively showed 30.1%, 5.9%, 32.5% and 10.3% binding to hemin-agarose, respectively. Again, this suggested that the kelch domain of K13 could confer the same efficiency of heme binding to KEAP1 as seen associated with K13/TrK13-WT protein. When soluble cellular extracts from stable transgenic Du145 cells ectopically expressing the K13-GFP, KEAP1 and their chimera proteins were used, we again found that K13-GFP, KEAP1, KEAP1^N^K13^KRP^-GFP and K13^N^KEAP1^KRP^-GFP bound hemin-agarose beads with efficiencies of 58.9%, 3.1%, 64.5% and 29.3%, respectively (**Fig. 5g**). Cumulatively, the results in **Fig. 5** strongly support that the kelch domain of K13 transfers robust heme binding capacity and heme-based stabilization to KEAP1. The N-terminal domains of K13 may also contribute a minor role consistent with our prior studies suggesting a subset of heme binding motifs are also detected outside of K13’s Kelch domain^24^.

### K13 and its chimeras confer sensitivity to heme-dependent ART-killing of Du145 cells

Since heme is implicated in the activation of ART, we next assessed the ART sensitivity of Du145 cells expressing K13-GFP and its chimeras (as well as GFP alone) in a standard killing MTT [3-(4,5-dimethylthiazol-2-yl)-2,5-diphenyltetrazolium bromide] assay (see *Materials and Methods*) comparing outcomes in HDM or media replenished with 5 µM heme. Du145 cells expressing control GFP showed an IC_50_ of 117.7 ± 0.9 µM in the absence of added heme (**Fig. 6a i**), thereby indicating that they are intrinsically highly resistant to ART. The addition of 5 µM heme minimally reduced the IC_50_ by 2.6%. Expression of KEAP1 also showed a small effect inducing approximately 6.1% reduction. However, K13-GFP and the KEAP1^N^K13^KRP^ cells had a greater effect, which translated into increasing heme-dependent sensitivity to ART by 18.2% and 23.7%, respectively (**Fig. 6a i-ii**). K13^N^KEAP1^KRP^-GFP had lesser effect showing ART sensitivity increase of 13.5%. The relative order of conferring increased ART sensitivity was KEAP1^N^K13^KRP^-GFP > K13-GFP > K13^N^KEAP1^KRP^-GFP > KEAP1 > GFP, which matched the heme binding properties reported for these transgenes in **Fig. 5**. Together, these data suggested that the presence of K13, particularly its kelch domain, conferred sensitivity to ART in the presence of 5 µM heme.

**Fig. 6.**
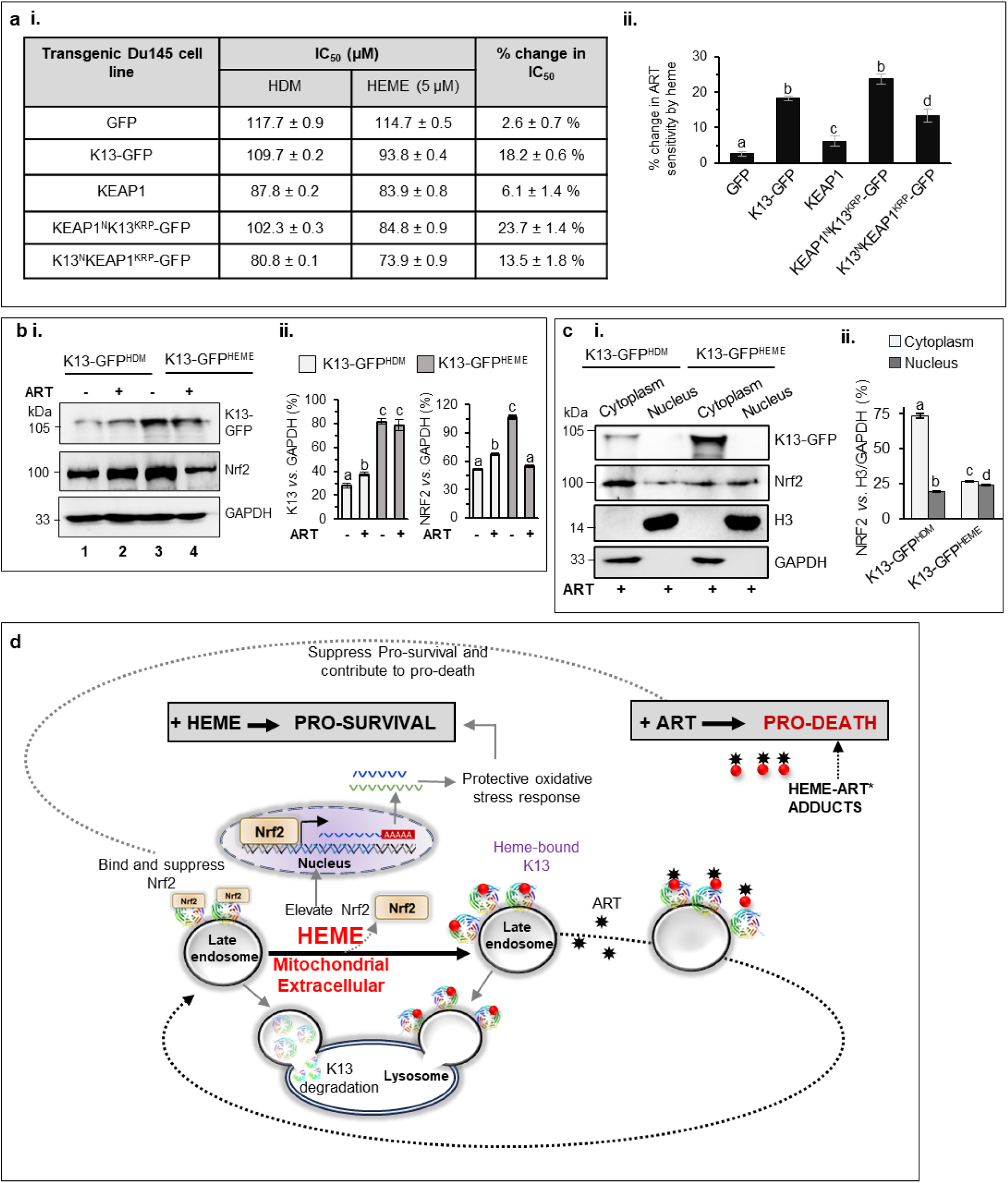
Dynamics of K13, heme and Nrf2 in a redox survival mechanism that promotes ART-induced death. **a.** IC_50_ values (µM, i) and change in percentage sensitivity to ART (right, ii) in respective transgenic Du145 cell lines on supplementation with 5 µM heme. **b.** Western blots (i) and quantitative densitometric analysis (versus GAPDH, ii) for K13-GFP (top) and Nrf2 (middle) in K13-GFP^HDM^ (lanes 1 and 2) and K13-GFP^HEME^ (lanes 3 and 4) cells in the absence (lanes 1 and 3) or presence (lanes 2 and 4) of ART. GAPDH (bottom) is the loading control. **c.** Western blots (i) and graphical densitometric quantitation (ii) for cytoplasmic and nuclear fractionation of K13-GFP and Nrf2 in K13-GFP^HDM^ and K13-GFP^HEME^ cells in the presence of ART. Molecular weight (in kDa) are as indicated. Error bars in a-c represent mean of three technical replicates ± SE. Statistical significance was calculated by one-way Anova. Distinct alphabets represent significant difference at p.adj ≤ 0.05. **d.** Model for a redox survival mechanism that promotes artemisinin-induced death. Dynamics of Nrf2 binding by free K13, its displacement by heme inducing (i) a redox survival response and (ii) K13-heme mediated activation of ART inducing oxidative pro-death response in Du145 cells.

Since the K13 also binds Nrf2 (which is antagonized by heme), we examined the effects of ART on the dynamics of K13 and Nrf2 in HDM as well as after heme supplementation. As shown in **Fig. 6b i-ii**, ART had no sizeable effect on K13-GFP levels in the presence or absence of heme. However, Nrf2 levels were reduced by 50% in the presence of both ART and heme. This reduction was due to the depletion of Nrf2 in the cytoplasm (**Fig. 6c i-ii**), suggesting it was through regulation by K13. These data support a model shown in **Fig. 6d**, where heme plays a major role in binding K13 to displace Nrf2, which in turn is targeted to the nucleus to induce a protective redox (or pro-survival) response. Heme bound to K13 activates ART to generate toxic heme-ART adducts. K13’s renewed ability to regulate Nrf2 suggests that these toxic heme-ART adducts do not inactivate K13. Rather, the adducts dislodge from K13 and disperse to mediate damage which occurs in absence of a protective redox response, resulting in cell death (or pro-death) responses. Thus, ART-induced killing reconstituted in Du145 cells may arise from the dual effect of inducing toxic ART-heme adducts as well as preventing a protective redox response to render an overall ‘pro-death response (**Fig. 6d**).

### ART sensitivity is increased proportional to the K13 intensity

Our findings in **Fig. 6** strongly supported that expression of K13 or chimeras, containing its kelch domain in presence of 5 µM heme, increased sensitivity to ART. We wanted to gain better understanding on whether increasing K13 at sub-micromolar heme concentration also played a role. Since our studies in **Fig. 4** established that ATRA increased levels of K13, we separately evaluated the effects of heme and ATRA. As shown in **Fig. 7a, b**, in cells expressing GFP, 5 µM heme increased sensitivity to ART by ∼ 6%, while ATRA had no effect. In K13-GFP cells, heme alone stimulated ART-sensitivity by 18%, ATRA by 25%, and the combination of two stimulated ART-sensitivity by 36% (**Fig. 7a, b**). These data strongly suggested that elevation in levels of K13-GFP by ATRA even under HDM conditions increased sensitivity to ART. But heme at 5 µM provided additional sensitization effect possibly by (i) increasing K13 levels (ii) providing substrate to form toxic heme-ART adducts, whose disassociation from K13 suppression of pro-survival responses (as hypothesized in **Fig. 6d**).

**Fig. 7.**
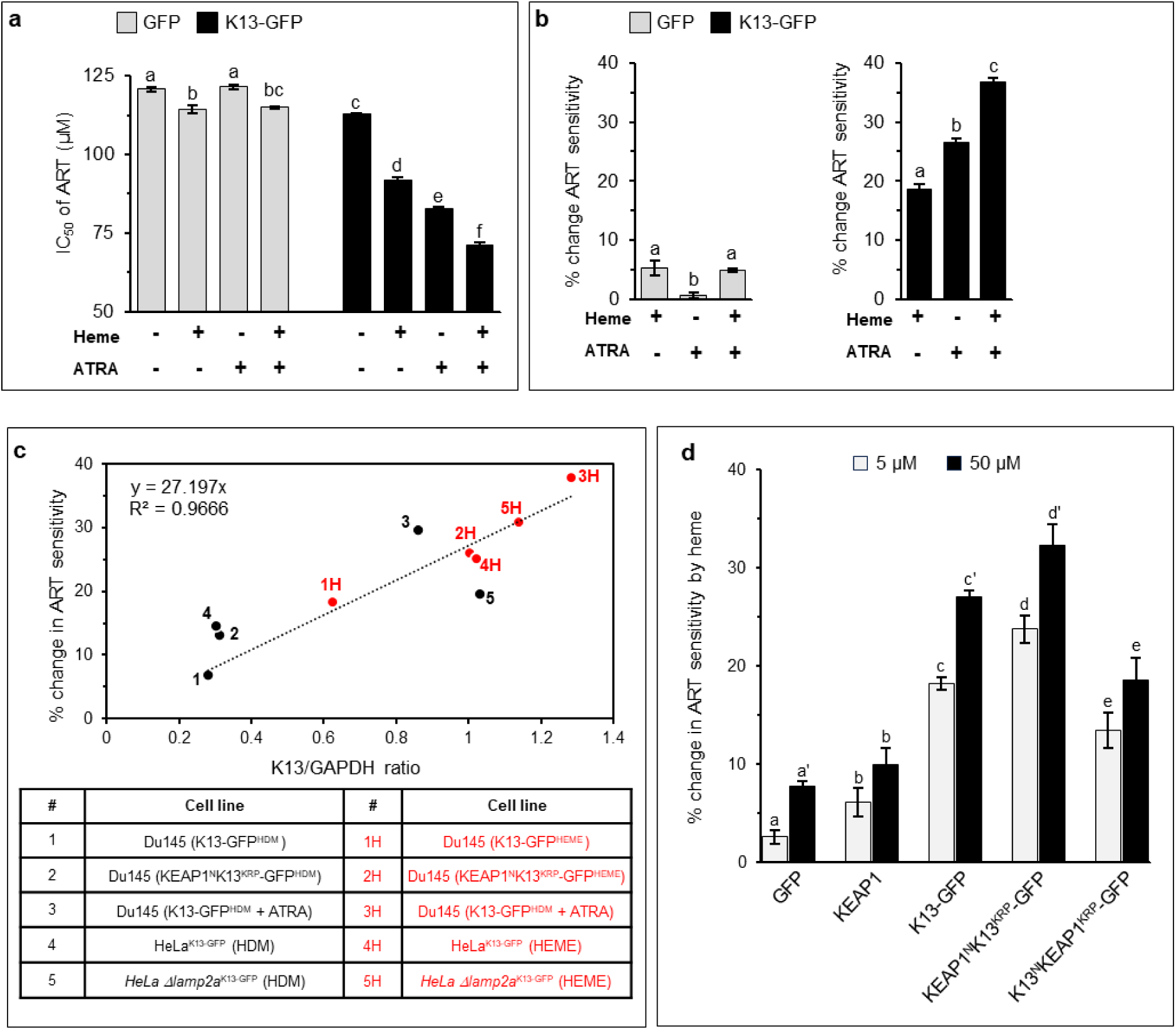
ART sensitivity is increased proportional to K13 intensity. **a.** IC_50_ values (µM) for GFP and K13-GFP cells in presence or absence of ATRA or heme (5 µM). **b.** Effect of heme (5 µM) and ATRA on percentage change in ART sensitivity in cells expressing GFP (grey bars) or K13-GFP (black bars). **c.** Graph showing the percentage change in ART sensitivity as a function of K13/GAPDH intensity ratio under HDM (solid black circles) or 5 µM heme-repleted (solid red circles) conditions in transgenic Du145 and HeLa cells. Black dotted trendline and R-squared values are indicated. Description of cells is shown at the bottom. Numbers only and numbers followed by initial ‘H’ indicate cells cultured under HDM or 5 µM heme-repleted conditions, respectively. **d.** Graph showing change in percentage sensitivity to ART at extracellular 5 µM (grey) and 50 µM (black) heme in the repleted media. Error bars in a, b and d represent mean of three technical replicates ± SE. Statistical significance was calculated by one-way Anova. Distinct alphabets represent significant difference at p.adj ≤ 0.05.

We were therefore interested in determining whether ART sensitivity was proportional to K13 intensity. To do so, we compared the sensitivity to killing by ART to levels of K13 (or its kelch chimera) normalized to GAPDH (a control cytoplasmic protein). As shown in **Fig. 7c**, we observed an overall linear relationship between increase in ART sensitivity and the K13/GAPDH (R^2^ value = 0.97). Notably, cell lines designated as 1, 2 and 4 reflected measurements generated in HDM, and all showed lower degree of ART sensitivity and K13/GAPDH levels than counterpart measurements generated in the presence of 5 µM heme. Also, the presence of ATRA increased K13 levels, thereby boosting K13/GAPDH ratio and correspondingly showed increased ART sensitivity of K13-GFP cells under HDM conditions (as cell line #3 in **Fig. 7c**). Moreover, combination of heme and ATRA in these cells (designated as 3H) further elevated K13/GAPDH ratio and showed highest sensitivity to ART (38%). Comparisons in HeLa cells revealed that knockout of LAMP2A, which increased K13/GAPDH by three-fold raised ART sensitivity by one-third. However, 5 µM heme further raised the K13/GAPDH ratio four-fold and doubled ART-sensitivity (compared to parental HeLa in HDM).

When we separately compared the effects of extracellular 5 µM *versus* 50 µM heme (**Fig. 7d**), the higher concentration of heme further augmented killing. In GFP cells, this could be easily interpretable as increasing intracellular heme alone from ∼400 to 750 nM directly sensitized ART-killing, increasing it from ∼2 to ∼6%. However, as we have previously shown for K13 kelch domain proteins, heme may also act by raising levels of the transgene alone. We therefore tracked transgene expression at 50 µM heme (**Fig. 7d**, black bars). This revealed that even at 50 µM, increasing K13-GFP and K13 chimeras stimulated ART-killing to much higher levels. Taken together, these data suggest that heme alone (at intracellular concentrations of 400 nM to 1 µM), is comparatively weak stimulator of ART’s killing activity but can be potentiated by expression of K13 and its appropriate chimeras.

## Discussion

K13 was identified as marker of ART-R more than a decade ago^6^. Our studies provide the first evidence that it acts as redox sensor regulated by the oxidant heme (summarized in **Fig. 8a**). This is also the first report that a small molecule like heme can bind a member of an evolutionary conserved kelch family (whilst multiple protein partners have been defined for kelch proteins^15,59–61^). Notably, K13-heme activates ARTs, suggesting the potent killing action of these drugs exploits a parasite protective redox response (**Fig. 8a**, cycle 1) Whether *Plasmodium* encodes for functional orthologue of Nrf2 (the transcription factor that induces the stress-induced protective response in Du145 cells) remains unknown. However, since heme is a strong oxidant^31^, upon its binding to K13, a protective redox response is likely elicited as a compensatory effect (expected for **cycles 2 and 3** shown in **Fig. 8a**). A redox stress response may be needed to protect the parasite against high levels of heme reported at all stages of parasite growth^62^.

**Fig. 8.**
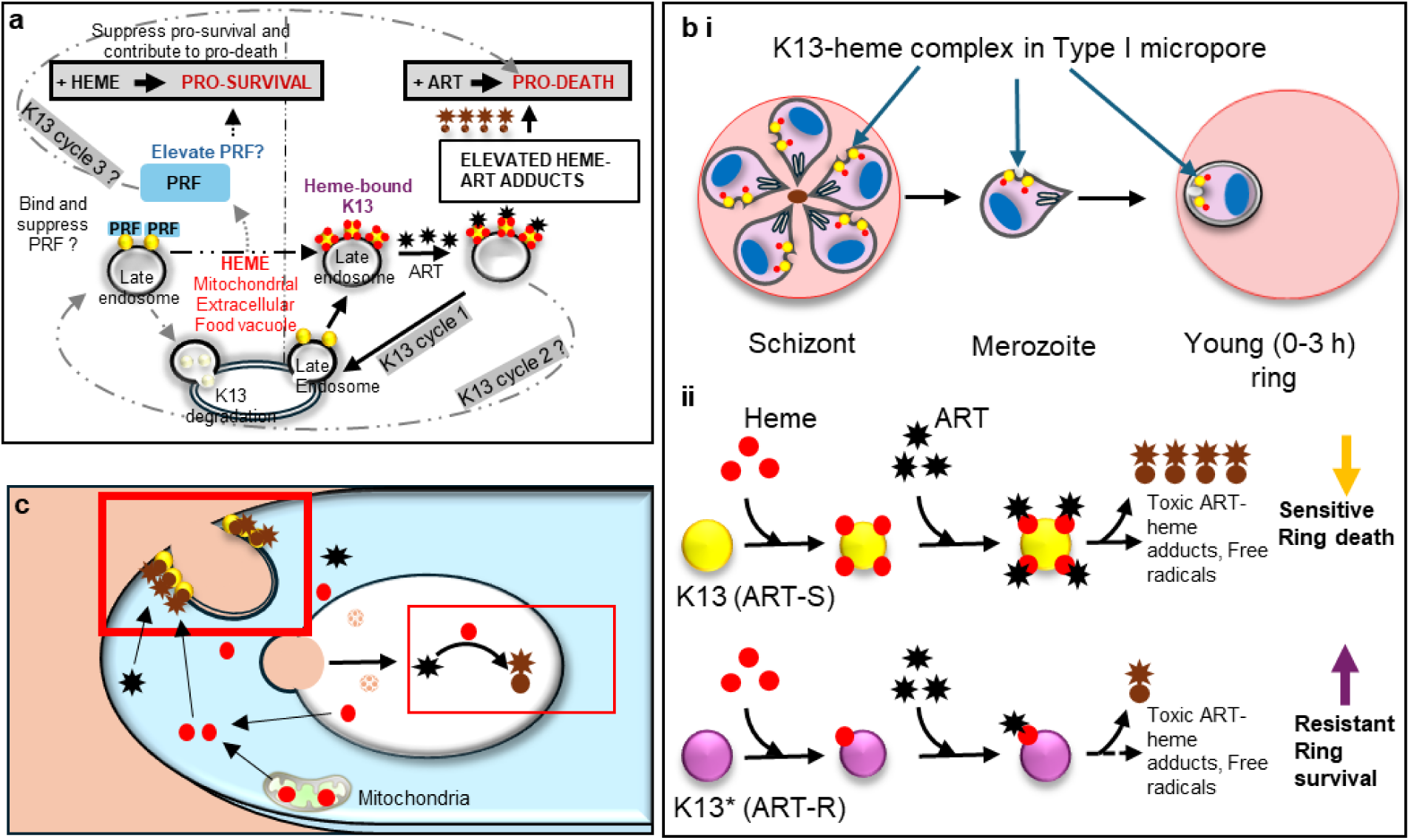
A new model for clinical ART-R, implicated by K13-heme redox survival responses and ART-induced death. **a-b.** ART induced pro-death mechanisms and their modulation in rings. Our primary conclusion is for **K13 in cycle 1** (bold lines, **a**), where K13 (solid yellow circle) binds heme (solid red circles, **a**) and becomes stabilized at cytoplasmic face of endosomal Type I micropore membranes of merozoites and rings (shown in panel **b**). For merozoites and rings, heme may come from mitochondrial and extracellular sources. K13-heme activates ART (solid black stars) to heme-ART adducts (dark brown hybrid star-circles), which dissociate from K13 (resulting in suppression of K13 redox sensing). Widespread toxicity of heme-ART adducts to cellular targets, triggers a pro-death response in the parasite. Heme-free K13 may re-cycle back to resume heme binding and re-initiate cycle 1 and reinforce death outcomes. **K13 cycle 2** may be initiated if a (yet hypothetical) parasite redox factor (PRF; solid blue rectangle) binds K13 and becomes suppressed until heme-binding dislodges the PRF to activate ART and pro-death effects, acting additively with cycle 1. **K13 cycle 3** may be initiated upon K13 displacement of PRF to trigger pro-survival responses and reduce pro-death effects of cycles 1 and 2**. b i**. Model proposing K13-heme complexes (solid yellow circles with solid red circles) assemble at Type I micropores of *P. falciparum* merozoites as they develop in schizonts. Released merozoites invade new red blood cells to form rings associated with Type I micropores and K13-heme complexes^71^. **b ii.** In wild type rings, K13-heme trigger ART (solid black stars)-induced oxidative damage radicals that destroy essential Type I micropores, resulting in ring-death. K13 ART-R mutants (solid purple circles) show reduced heme binding, oxidative damage enabling ring survival and ART-R. **c.** K13-heme is also catalytic for ART activation in trophozoites. K13 (solid yellow circles) concentrates at cytostome structures that enable hemoglobin digestion and liberation of free heme (solid red circles) in the food vacuole. Existing models propose that free heme activates ART (solid black stars) to toxic heme-ART adducts (dark brown hybrid star-circle) in the food vacuole (see red box). We propose that heme from the food vacuole and other sources (like the mitochondria) transported to the parasite bind K13 concentrated at the cytostome, which greatly stimulates formation of activated ART, and toxic heme-ART adducts (dark brown hybrid star-circle in bold red box) compared to encounters of ART with free heme in the food vacuole, thus explaining why K13 mutants do not confer ART-R at these stages.

The models in **Fig. 8a, b** maybe particularly informative for clinically reported ART-induced ring-stage death^63,64^ based on the following considerations. Young rings (0-3 h) lack a digestive food vacuole and therefore depend on heme from biosynthetic sources, analogous to Du145 cells. Similarly in *P. falciparum* rings, K13-heme triggers a pro-survival response exploited by ART to spring a pro-death response of oxidizing ART-heme adducts that are no longer redox active but can kill the ring parasite (**Fig. 8b i**), and this function is compromised in ART-R (**Fig. 8b ii**). To date, K13 function has been best understood in context of regulation of uptake of hemoglobin into the food vacuole at the trophozoite stage. However, it is well documented that K13 is expressed in daughter merozoites that assemble when trophozoites mature to the segmented schizont stage. K13 is also detected from the earliest stages of ring infection (mode in **Fig. 8b ii**)^59,65,66^. However, K13’s precise location is not well mapped in *P. falciparum* merozoites and rings. But in the related apicomplexan *Toxoplasma gondii*, a K13 orthologue was localized to endosomal structures called Type I micropores and shown to regulate plasma membrane endocytosis^67,68^. Since all apicomplexans (including *Plasmodium*) contain a ‘Type I’ micropore in their ‘apical’ stages^69,70^, by virtue of evolutionary conservation of mechanisms of apical organelles, we have proposed that in merozoites and rings K13 localizes to *P. falciparum* Type I micropores and regulates essential plasma membrane endocytic functions of these parasite stages ^71^ (**Fig. 8b i**). Indeed the *P. falciparum* food vacuole where K13 is abundant develops from a Type II micropore, which is also consistent with a conserved micropore function for K13.

The Du145 cell model enabled assessment of the effects of K13 binding to heme from nanomolar to sub-micromolar range. The intrinsic concentration of heme in Du145 cells is 60 nM. Supplementing with extracellular heme at 5 µM and 50 µM resulted in maximal intracellular concentrations of 0.45 µM and 0.75 µM, which approach but are nonetheless lower than the published physiological range (1-5 µM) reported for *Plasmodium*^40,41^. Whether kelch proteins of other apicomplexans bind heme has yet to be investigated. Moreover, although the kelch domain of K13 is necessary and sufficient to confer heme binding property to its closest mammalian orthologue KEAP1, whether and how heme is released from K13 under physiological conditions is not known, since it cannot be ubiquitinylated and degraded, which is a hallmark of protein substrates and their cellular regulation by kelch proteins. But our studies suggest that when K13-bound heme activates ART, the kelch structure may protect against self-oxidation by heme-ART adducts^24^. This is consistent with prior studies showing that alkyne-ART analog despite being highly active against a wide range of malarial cellular targets failed to alkylate K13^72,73^. Heme-ART adducts show low affinity for recombinant K13^24^, suggesting a mechanism for their release and a new model where a single molecule of K13 may catalyze multiple cycles of heme-ART toxic adduct formation and oxidative killing (**Fig. 8a**; K13 cycle 1).

That we find that intracellular heme concentrations of 400-750 nM alone yield low sensitization to ART, counters prevailing models that suggest that heme alone is sufficient to activate a low level of ART at the trophozoite stages (**Fig. 8c**). Notably, our data show that even at the lowest intensities, K13 (expected not to exceed nanomolar levels) was more effective than heme alone and in addition, the extent of sensitization was directly proportional to the level of K13. This is consistent with our findings that raising K13 (by ATRA treatment) in HDM increased ART-sensitivity more than an eight-fold in free heme concentrations of 60 nM. These data strongly support the idea that K13 acts as ‘catalyst’ to facilitate assembly of activated ART-heme complexes, and once the activated toxic adducts are released, K13 may facilitate successive rounds of ART-based killing. We therefore propose that even at the trophozoite stage, rather than heme alone, it is the K13-bound heme which effects sensitization to ART (**Fig. 8c**). These stages are known to be far more sensitive to ART than rings, and trophozoites expressing K13 mutations of ART-R are highly susceptible to killing by ART, which is consistent with their higher levels of K13.

K13-heme concentration at late endosomes in Du145 cells suggests that in addition to triggering a pro-survival response, heme binding to K13 may stabilize endosomal location in both rings and trophozoites and possibly create concentrated cellular foci of K13 function as well as efficiently activate ART at all stages of parasite growth. That heme binding, endosomal concentration and ART-based killing are not properties of mammalian KEAP1, suggests they are parasite evolutionary adaptations conferred by the K13 kelch domain. Yet, orthologues of LAMP2A (a receptor for K13 in Du145 and HeLa cells that controls its autophagic degradation), have not been reported in *P. falciparum* or *T. gondii.* Nonetheless, autophagic systems exist in both parasites^74,75^ and additional studies are needed to determine their influence on heme-dependent K13 stability, as well as endosomal degradation and turnover of K13 and its orthologues across apicomplexans. Another key priority is identification of K13-coupled parasite redox factors that remain unidentified because a lack of conservation at the level of primary sequences prevents easy prediction from nucleotide sequence comparisons. While our studies establish that K13 is an important determinant in ART sensitization, other redox and oxidative factors may also play a role. Our data suggest Du145 may continue to provide a powerful model system for future functional screens of endosomal effectors as well as novel plasmodial pro-oxidant responses, regulated by both heme and K13 and targeted in killing by ART. One generalized lesson already learned may be that suppression of redox sensing may sensitize a wider range of infectious agents as well as cancers to killing by ARTs.

## Materials and methods

### Generation of plasmid constructs

The pPR-IBA101 (TrK13-WT) plasmid was generated as described in earlier studies^24,47^. For the generation of pPR-IBA101 (KEAP1), the full length (1875 base pairs) transcript of *keap1* was amplified from cDNA library prepared from human cell line (HEK293T) using the primer pairs (Hs-Keap1-BsaI-Fwd (5’-CAAATGGGAGACCTTATGCAGCCAGATCCCAGGC-3’) and Hs-Keap1-BsaI-Rev (5’-CCAAGCGCTGAGACCAGCAGCACAGGTACAGTTCTGCTGGTCAATCTG-3’). The PCR product was digested with BsaI and cloned at corresponding sites in similarly digested pPR-IBA101. For the generation of pPR-IBA101 (KEAP1^N^K13^KRP^), nucleotide sequences coding for the N-terminus region of KEAP1 was amplified from pPR-IBA101 (KEAP1) plasmid using (Hs-Keap1-BsaI-Fwd and KEAP1toK13-KRP-Rev (5’-AAATCCACCGGCAGCAGCGCGGCCCACCTTGGG-3’) primers. Nucleotide sequence encoding for the KRP region of K13 was amplified using the pPR-IBA101 (TrK13-WT) plasmid as template and KEAP1toK13-KRP-Fwd (5’-GTGGGCCGCGCTGCTGCCGGTGGATTTGATGGTGTAGAATATTTAAATTCG-3’) and PfKelch13-BsaI-Rev (5’-GGAGGACAAAATGGCAATGTTCTAAATTCATGTGGATTCTTTTCACCAGATACAAATGAATGGC-3’) primer pairs. The two PCR products were used as templates to reconstitute KEAP1^N^K13^KRP^ by overlapping PCR, which was then digested with BsaI and cloned into pPR-IBA101. Similarly, to generate pPR-IBA101 (K13^N^KEAP1^KRP^), nucleotide sequences encoding for the N-terminus of K13 was amplified from pPR-IBA101 (TrK13-WT) plasmid as template and PfKelch13-BsaI-Fwd (5’-CAAATGGGAGACCTTATGGAAGGAGAAAAAGTAAAAACAAAAGCAAATAGTATCTCG-3’) and K13toKEAP1-KRP-Rev (5’-GTAGATCAGGGCAGCAGCTATACAAAATACTAATGGGAATGGTAAAAATTTAATACC-3’) primers. Overlapping PCR, using the above two PCR products as templates, generated K13^N^KEAP1^KRP^ which then introduced to pPR-IBA101 after digestion with BsaI.

For generating recombinant Nrf2-6×his, the full length (1626 bp) transcript was amplified from cDNA library prepared from human cell line (Du145) using the primer pairs HsNrf2-XbaIFwd (5’-CACGTGTCTAGAATGATGGACTTGGAGCTGCCG-3’) and HsNrf2ns-XhoIRev (5’-CCTAGGCTCGAGGTTTTTCTTAACATCTGGCTTCTTACTTTTGGG-3’) and the PCR product was then digested with XbaI-XhoI and clone into similarly digested pET303-ct-his plasmid (ThermoFisher, USA) for in-frame fusion with C-terminus 6×his tag. To generate GFP expressing constructs for mammalian cells, pAcGFP1-C1 (Addgene, USA) was used and AgeI and NheI sites considered for inserting the gene of interest. To generate pAcGFP1-C1 (K13-GFP), pAcGFP1-C1 (KEAP1^N^K13^KRP^) and pAcGFP1-C1 (K13^N^KEAP1^KRP^) plasmids, their pPR-IBA101 plasmids were used as corresponding templates. For K13-GFP, the primers pairs were PfKelch13-NheI-Fwd (5’-AAGCTTGCTAGCATGAACAGCGAGGTGTGTTCGCG-3’) and PfKelch13-AgeI-Rev (5’-AGCGCTACCGGTAGCAGCCAAATTTGCCAAGAGAATGCCGTG-3’), for generating KEAP1^N^K13^KRP^GFP, the primer pairs were KEAP1-NheI-Fwd (5’-AGATCTGCTAGCATGCAGCCAGATCCCAGGC-3’) and PfKelch13-AgeI-Rev (5’-AGCGCTACCGGTAGCAGCCAAATTTGCCAAGAGAATGCCGTG-3’); and for generating K13^N^KEAP1^KRP^, PfKelch13-NheI-Fwd (5’-AAGCTTGCTAGCATGAACAGCGAGGTGTGTTCGCG-3’) and KEAP1-AAA-AgeIRev (5’-GGATCCACCGGTAGCTGCTGCACAGGTACAGTTCTGCTGGTCAATC-3’) primers were used. To generate the KEAP1 construct, pPR-IBA101 (KEAP1) plasmid was used as the template to amplify nucleotides encoding for KEAP1-strep using KEAP1-NheI-Fwd and Strep-XhoI-Rev (5’-GGATCCCTCGAGCTTTCTCGAACTGCGGGTGGCTCCAA-3’). The PCR amplified product was subjected to NheI-XhoI digestion and inserted into similarly digested pAcGFP1-C1.

All constructs generated for this study were confirmed by Sanger’s sequencing prior to further experiments.

### Cell culture, generation of transgenic lines and their treatments

The Du145 cell line was the kind gift from Prof. Rana Pratap Singh, SLS, JNU, India. The parental HeLa (ab255928) and HeLa-Δ*lamp2a* (ab255402) were procured from Abcam, USA and cultured according to the manufacturer’s instructions in complete media RPMI (Gibco, USA). Cells were transfected using Lipofectamine 3000 (Invitrogen, USA) following the manufacturer protocol. Stable transgenic lines were generated by G418 (0.5 mg/ml, Sigma, USA) selection. For the heme depleted (HDM) conditions, the cells were grown in RPMI media supplemented with heme-depleted FBS and succinyl acetone (0.5 mM, Caymanchem, USA) for 48 h. For heme-replenished media 5 μM hemin chloride (Merck, USA) was added to the HDM. To assess *in vivo* stability of K13 protein upon heme binding, the cells were treated with heme analogs PPIX (50 μM, Merck, USA) or Zn-PPIX (50 μM, Merck, USA) for 48 h. Cells were also subjected to treatment either with for 48 h with ART (50 μM, Caymanchem, USA) in presence or absence of heme.

### Preparation of heme-depleted FBS

Commercially purchased FBS (Gibco, USA) was incubated with ascorbic acid (10 mM, SRL, India) at 37°C in a shaking incubator at 200 rpm for overnight. Spectral scans of the pre- and post-treated FBS measured to check the presence or absence of soret peak at 410 nm, respectively using JASCO UV660 spectrophotometer. Finally, the ascorbate-treated FBS was dialyzed against PBS, pH 7.4, with at least three periodic changes for 24 h at 4°C. Dialyzed FBS was filter-sterilized using 0.2 μm polyether sulfone filter (Sartorius, USA).

### Assessment of cellular viability

Cell viability was assessed using 3-(4,5-dimethylthiazol-2-yl)-2,5-diphenyltetrazolium bromide (MTT, Merck, USA)-based colorimetric assay. Cells at 50-60% confluency and within 72 h incubation in the 96-well cell culture plate were only considered. Following treatment, the cells incubated with 20 µl of 5 mg/ml MTT solution for 2h at 37° in the dark. Finally, the culture media was discarded, and formazan crystal was dissolved with 200 µl of DMSO, and absorbance was recorded at 570 nm using a microplate reader (Thermo Fisher Scientific, USA). Cell viability was calculated and expressed as percentage in comparison to the untreated controls. For MTT assay, the viability was calculated for six different concentrations (0, 40, 80, 120, 160 and 200 µM) in three replicates to generate the IC_50_ values for ART.

### Estimation of total cellular heme content

Cells were grown up to 90-95% confluency and dislodged from the plate by treatment with trypsin and were washed twice with saline PBS. Cell count was recorded prior to lysis using hemocytometer. Cells were then solubilized in five volumes of RIPA buffer for 2 h at 4°C. After a brief centrifugation, the supernatant was diluted ten times and subjected for spectrophotometric scanning at 410 nm. The absorbance was plotted against the standard curve of hemin chloride to calculate the total cellular heme content. The standard curve has been generated using gradually increasing concentration (0, 0.05, 0.1, 0.5, 1 and 10 µM) of hemin chloride.

### Nuclear and cytoplasmic fractionation of transgenic cells

Cellular fractionation into nuclear and cytoplasmic fractions was performed following previously described protocol^76^. Briefly, the cells from the 60 mm culture plates were scrapped and washed twice with ice-cold phosphate buffered saline (PBS) pH 7.4. The cell pellets were washed, resuspended in 250 μl of ice-cold 0.2% (v/v) NP40 (HiMedia Labs, India), minced with micro-pestle, vortexed and kept on rocker shaker for 15 mins. The resuspended pellets were then centrifuged at 10,000 rpm for 1 min, and the supernatant was collected as the cytoplasmic fraction. The pellet was then resuspended in 1 ml ice-cold 0.2% (v/v) NP40 to remove remnant cytoplasmic debris and centrifuged at same speed. Finally, the pellet was resuspended in 200 μl ice-cold PBS and designated as the nuclear fraction. The nuclear fraction was also briefly sonicated using micro pulser (Sonics, USA).

### Indirect immunofluorescence assay (IFA)

IFA was performed using parental or transfected cell lines. Briefly, cells were initially grown on poly-L-lysine coated coverslips and fixed using 4% (w/v) paraformaldehyde/ 0.0075% (v/v) glutaraldehyde followed by permeabilization with 0.1% (v/v) Triton X-100 and neutralized with 50 mM NH_4_Cl. Blocking was done using 3% (w/v) BSA. The coverslips were incubated with respective primary antibodies (10 μg/ml) at 4°C for overnight followed by incubation in FITC- or TRITC/ Rhodamine-conjugated secondary IgG antibodies (1:200; MP biomedicals, USA). Cellular nuclei were stained with 5 μg/ml Hoechst 33342 (Molecular Probes) and slides were mounted with DABCO. The slides were finally visualized by SP8 confocal laser scanning microscope (Leica, Germany). Fluorescence intensity was calculated using Image-J software (https://imagej.net/ij/) software by considering 3 biological replicates for each set and each replicate containing 30 individual cells. The RGB intensity plot and Person’s correlation coefficient (PCC) was generated from the Image-J graphics and JaCop, respectively.

### SDS-PAGE and western blotting

The induced bacterial cell pellet was directly solubilized in Laemmli’s sample buffer and heat denatured for 15 min at 95°C. For mammalian cells, following scrapping from plates, cells were washed twice with saline PBS and solubilized in five volumes of RIPA buffer for 2 h at 4°C. After a brief centrifugation, the supernatant was considered as whole protein pool and boiled for 15 min at 95°C using Laemmli’s sample buffer and resolved by 8-10% SDS-PAGE followed by Coomassie or silver staining. For subsequent western blots, resolved proteins were transferred into nitrocellulose membrane (MDI, USA) followed by blocking with ice cold 5% (w/v) fat-free skim milk (SRL, India) for 2 h at RT. The membranes were incubated with specific primary antibodies (1:3000 to 1:5000 dilution) for overnight at 4°C on a platform shaker followed by subsequent washing and incubation with respective HRP-conjugated secondary antibodies (1:7000 to 1:10000 dilution). Blots were finally developed by ECL assay and detected using gel imager (Bio-Rad, USA). The primary antibodies are used in this study were: anti-GFP (Invitrogen, USA; Cloud clone, China), anti-UBQ, anti-Lamp2 and anti-GAPDH (Cloud clone, China), anti-k13 (Custom generated, Biobharati Life Sciences^47^), anti-Histone 3 (kind gift from Prof. Suman Kumar Dhar, SCMM, JNU, India), anti-Nrf2, anti-KEAP1, anti-HO1, anti-NQO1, anti-MCM3, anti-DPP3 and anti-TRAF2 (ABclonal, Woburn), anti-His and streptavidin-HRP (BioBharati Life Sciences, India), and anti-HA. The subsequent HRP conjugated secondary antibodies used in this study were goat anti-rabbit (Bio-Rad, USA), goat anti-mouse, goat anti-rat (MP biomedicals, USA). Band intensity was calculated from Image-J (https://imagej.net/ij/) software by considering 3 individual blots for each data set.

### Purification of recombinant protein and *in vitro* trypsin digestion

Recombinant Trk13-WT, KEAP1, K13^N^KEAP1^KRP^and KEAP1^N^K13^KRP^ proteins as C-terminus fusion with strep tag and Nrf2-6×his were purified from isopropyl-1-thio-β-D-galactopyranoside (IPTG, SRL, India) and arabinose (0.2%, SRL, India) induced bacterial pellet. Briefly, the induced bacterial pellet was lysed using lysozyme (SRL, India), sonicated and washed thrice with 2% (w/v) sodium deoxycholate (SRL, India) to purify inclusion bodies (IBs). The IBs were dissolved into 2 M urea, 50 mM Tris-HCl buffer pH 12.5 and then dialyzed against buffer W (50 mM Tris-HCl, 150 mM NaCl, pH 8). Dialyzed solution was kept for binding to StrepTactin XT Superflow resin (IBA, Germany) for Trk13-WT, KEAP1, K13^N^KEAP1^KRP^and KEAP1^N^K13^KRP^ and Ni-NTA resin (Thermo, USA) for Nrf2-6×his. Purified proteins were dialyzed overnight at 4°C against buffer W with at least three periodic changes. Purity of each protein was checked by SDS-PAGE and Coomassie staining and further confirmed by western blotting using commercial antibodies against the 6×his tag, strep tag (Biobharati Life Sciences) or custom-generated anti-K13.

For *in vitro* trypsin digestion, 10 µg of recombinant TrK13-WT or commercially procured BSA (A3294, Sigma, USA) was incubated with or without equimolar concentrations of heme in buffer W for 1 h. Samples were desalted as described in our earlier study^24^, and treated with 0.5 µg trypsin (T6567, Sigma-Aldrich) for different incubation times. Following digestions, Laemmli’s sample buffer was immediately added to each sample, boiled for 15 min at 95°C and processed for SDS-PAGE and Coomassie staining.

### *In vitro* protein-protein interactions

The purified recombinant Nrf2-6×his (10 µM) was incubated with Ni-NTA beads overnight and washed thrice with buffer W containing 30 mM imidazole. The Ni-NTA bound recombinant Nrf2-6×his was then incubated with purified recombinant TrK13-WT (10 µM) in buffer W for 4 h at RT. The beads were thoroughly washed with buffer W, and Ni-NTA-bound proteins were resolved by SDS-PAGE followed by Coomassie staining or western blot using antibodies against the 6×his tag, strep-tag (Biobharati Life Sciences) or custom-generated anti-K13. For experiments involving the interactions between heme-bound or heme-free recombinant TrK13-WT and Nrf2-6×his, the TrK13-WT was preincubated with heme (0 or 10 µM) at RT for 1 h at room temperature (RT) and desalted using PD Spintrap™ G-25 columns (Cytiva) to remove excess unbound heme.

### Hemin agarose binding

Affinity-purified recombinant Trk13-WT, KEAP1, K13^N^KEAP1^KRP^and KEAP1^N^K13^KRP^ proteins (10 µM) were incubated with 50 µl suspensions of 50% v/v hemin-agarose (Merck, USA) beads for 1 h at RT in buffer W, pH 8, under rotating conditions, followed by five washes with ten volumes of buffer W to remove unbound proteins. The hemin-agarose beads were then solubilized in Laemmli’s sample buffer, resolved by SDS-PAGE, and visualized by Coomassie staining. For assessing hemin-agarose binding from transgenic cellular lysates, the total cell pellet was resuspended in 5 volumes of ice-cold RIPA buffer complemented with 1X protease inhibitor cocktail (cOmplete, Roche) and incubated at rotating wheel for 1 h. After centrifugation (15 mins, 21,100 × g, 4°C), the supernatant containing soluble proteins fractions was incubated with hemin agarose beads for 4 h. Unbound proteins in the lysates were washed with ten volumes of RIPA buffer and bound fractions were then solubilized in Laemmli’s sample buffer and subjected to western blotting using anti-strep tag antibody (Biobharati Life Sciences).

### Co-immunoprecipitation

Protein A beads (IBA Life Sciences, Germany) were incubated with 10 µg of respective primary antibodies for 4 h at 4 °C under rotating conditions and unbound antibodies were removed by extensive washes with pre-cooled PBS. Mammalian cells were harvested, lysed with five volumes of RIPA buffer supplemented with 1 mM PMSF (Thermo, USA) and protease inhibitor cocktail (Roche) and centrifuged at 10,000 × g for 15 min at 4°C to remove the cell debris. Clarified lysates were then incubated with antibody-coated protein A beads (30-50 µl, 50% v/v resuspension) for overnight at 4°C under rotating conditions. Following binding, unbound lysates were removed and the beads washed twice with RIPA buffer and twice with ice-cold saline PBS. Beads were boiled with Laemmli’s sample buffer for 15 min at 95°C and processed for western blotting.

### Total RNA, cDNA synthesis, and real time RT qPCR

The total RNA was isolated by Trizol reagent (Invitrogen, USA). Briefly, cell pellets were washed twice with PBS and homogenized using 500 µl of Trizol reagent. The supernatants were isolated after a brief centrifugation, mixed with 300 µl chloroform and incubated at RT. After centrifugation at 13,000 × g for 15 min at 4°C, the upper aqueous phase was removed and 500 µl isopropanol added. Following incubation for 30 min at RT, the tubes were centrifuged at 13,000 × g for 15 min at 4°C. The pellet was washed twice with 70% ethanol and finally dissolved in TE. Samples were treated with DNase I enzyme (Thermo scientific, USA) to remove genomic DNA contamination. The quantity and quality of RNA was estimated using Bio spectrophotometer (Eppendorf, Germany) and agarose-formamide gel electrophoresis. Approximately 1 μg of total RNA was used for cDNA synthesis using PrimeScript 1st strand cDNA Synthesis Kit (Takara-Bio, Japan) following manufacturer’s protocol. For real time RT-qPCR analyses, 2 μl of ten-fold diluted stock solution of cDNA samples was used. To perform RT-qPCR analyses, primers were designed based on the coding sequence of specific genes using Oligo Analyzer software (https://eurofinsgenomics.eu/en/ecom/tools/oligo-analysis/). Primers pairs were HsHO1_RT_Fwd (5’-ATGACACCAAGGACCAGAGC-3’) and HsHO1_RT_Rev (5’-GCATAAAGCCCTACAGCAACT-3’); HsNQO1_RT_Fwd (5’-CCTTCCGGAGTAAGAAGGCAG-3’) and HsNQO1_RT_Rev (5’-TCCAGGCGTTTCTTCCATCC-3’); HsTXNRD1_RT_Fwd (5’-GGAACTAGATGGGGTCTCGG-3’) and HsTXNRD1_RT_Rev (5’-TCTTGCAGGGCTTGTCCTAA-3’). RT-qPCR analyses were performed by using HOTFIRE Pol EvaGreen qPCR Mix Plus (ROX) (Solis BioDyne, USA) and Step one plus system (Applied biosystem, USA). To confirm the absence of any primer dimers in the amplified products, a standard melting curve analysis was conducted. Primers for *gapdh* included HsGAPDH_RT_Fwd (5’-CAGGAGGCATTGCTGATGAT-3’) and HsGAPDH_RT_Rev (5’-GAAGGCTGGGGCTCATTT-3’) as reference. Relative fold change was calculated by 2^−ΔΔCt^ method.

### Statistical validation

One-way ANOVA was carried out to test significance in difference considering at least three biological replicates using GraphPad prism 10. The adjusted p values (p.adj ≤ 0.001) were obtained via Tukey’s HSD.

## Author contributions

Conceptualization Ideas: S.D., K.H., S.B.^1^ and S.B.^2^.

Methodology: S.D., M.F.

Validation: S.D.

Investigation: S.D.

Resources: S.B.^1^, S.B.^2^, K.H.

Writing: S.D., K.H., S.B.^1^

Supervision: K.H., S.B.^1^

Project Administration: S.B.^1^

Funding acquisition: S.B.^1^

## Acknowledgements

This work was supported by research grants from the Indian Council of Medical Research (ICMR), (58/25/2020/PHA/BMS) and Indian Ministry of Human Resource Development (MHRD) Scheme for Translational and Advanced Research in Sciences (STARS) (MoE-STARS/STARS-2/2023-0111) from the Government of India (SB). SD is a recipient of Department of Biotechnology Research Associate fellowship (DBTRA/2023 24/N/JNU/118) from the Government of India.

## Data availability

The authors declare that the source data supporting the findings of this study are available within the paper and the associated supplementary information files (Supplementary Figures 1-4).

## Materials & Correspondence

Correspondence and material requests should be addressed to Souvik Bhattacharjee

