## Supplementary data for "A *Plasmodium falciparum* redox survival mechanism licenses killing by artemisinins"

**A *Plasmodium falciparum* molecular mechanism of heme binding and sensitivity to artemisinins.**

**Supplementary data**

Supplementary data include four supplementary figures 1-4.

**Supplementary figures and legends**

### Supplementary Fig. 1

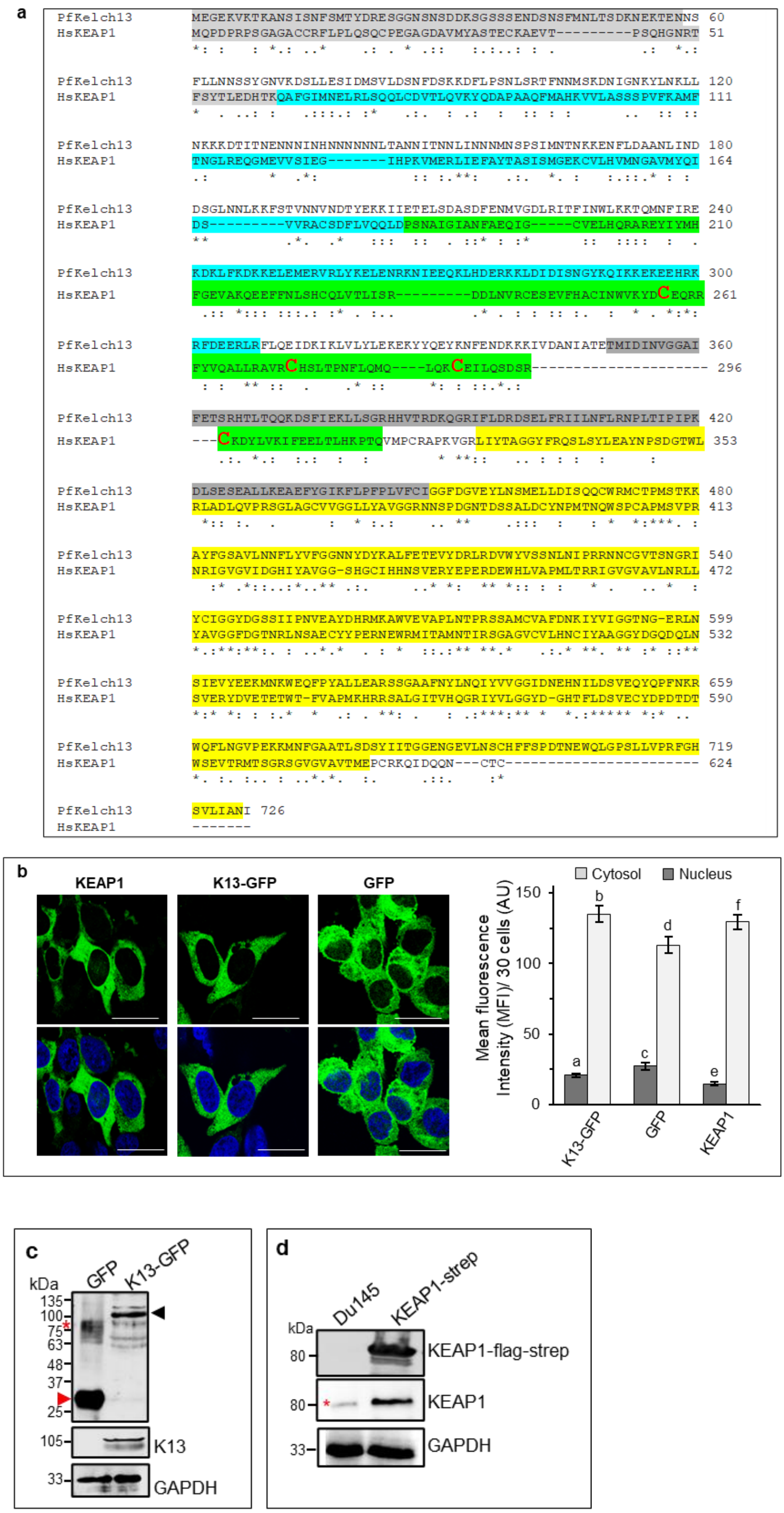

**Supplementary Fig. 1. Comparative evaluation of plasmodial K13 and human KEAP1. a.** Clustal Omega sequence alignment between K13 and KEAP1 indicating the N-terminus (light grey), coiled-coil (CCD; cyan), BTB/POZ (dark grey), intervening region (IVR; green) and kelch (yellow) regions. The four cysteine residues (Cys257, Cys273, Cys288 and Cys297) in KEAP1 implicated in KEAP1-dependent ubiquitination of Nrf2 and KEAP1-mediated repression of Nrf2 activity are shown in larger font and red color. **b.** Fluorescence micrographs of Du145 cells expressing the following transgenes (green): KEAP1-flag-strep (left), K13-GFP (middle) and GFP (right). Lower row shows associated nuclear staining with Hoechst 33342 (blue). Scale bar, 5 μm. Corresponding right hand side graphs indicate quantitative mean fluorescence intensity (MFI) values from 30 independent cells. Graph in b represent mean from three biological replicates ± SE. Statistical significance calculated by one-way Anova (Tukey's multiple comparisons test). Distinct alphabets represent significant difference at $p.adj \leq 0.05$. **c.** Expression of K13-GFP (black arrowhead) and GFP (red arrowhead) in transgenic Du145 cells, detected using anti-GFP (top) and anti-K13 (middle) antibodies. The loading control was GAPDH (bottom). Red asterisk marks non-specific smearing. **d.** Expression of KEAP1-flag-strep in transgenic Du145 cells, as detected by anti-strep (top) and anti-KEAP1 (middle) antibodies. Anti-KEAP1 antibodies also detected low levels of native KEAP1 protein (black asterisk) in parental Du145 cells. The loading control was GAPDH (bottom). Molecular weight standards (in kDa) are as indicated.

**Supplementary Fig. 2**

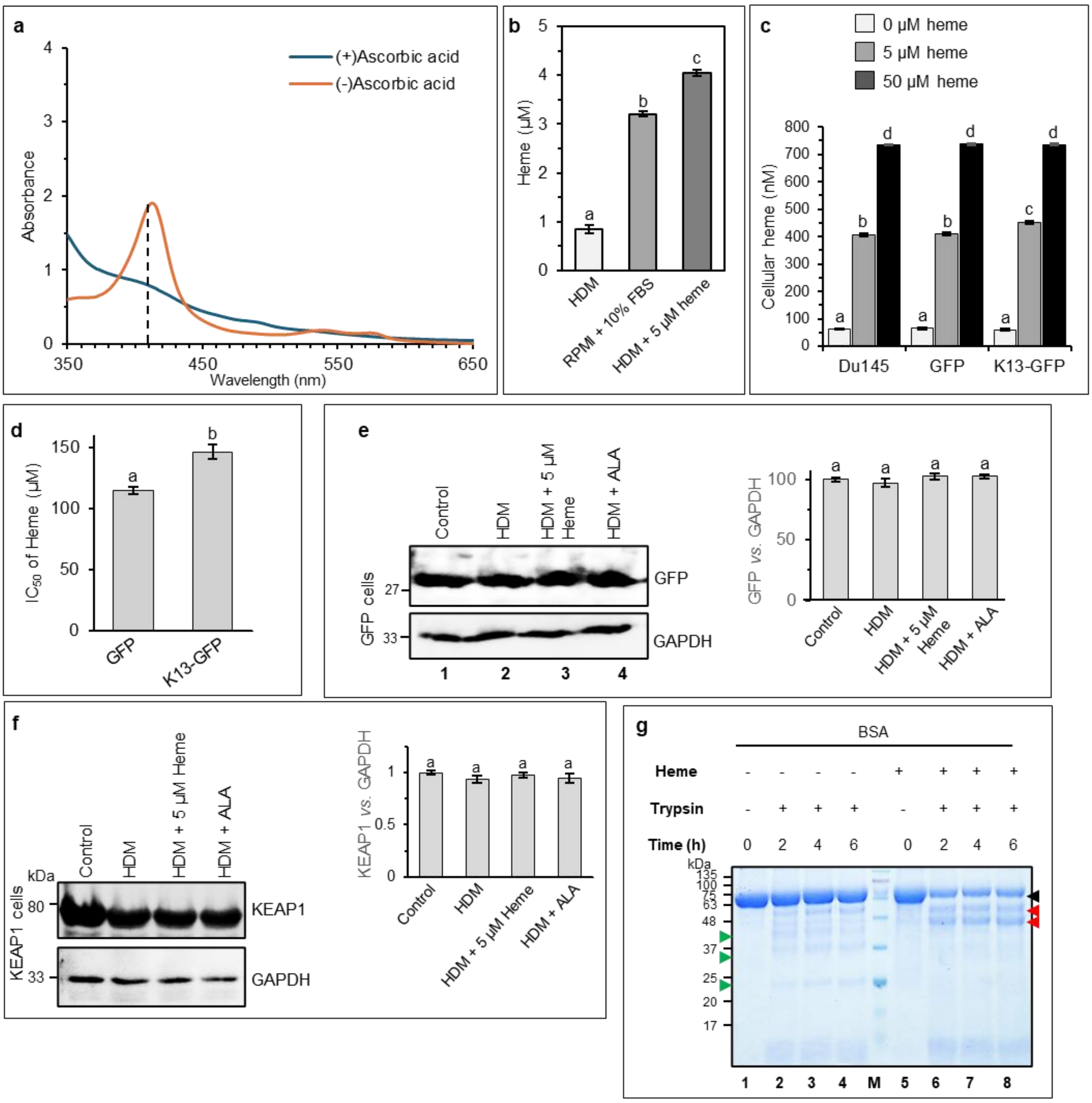


**Supplementary Fig. 2. Heme-dependent properties and effects characterized in Du145 cells as well as in *in vitro* measurements. a.** Spectrophotometric scans of commercially procured FBS (red) indicating a heme soret peak at 410 nm (dotted black line). This peak decreased when FBS was treated with 10 mM ascorbic acid (blue) indicating that it resulted in quantitative depletion of heme (showing potential to develop a method to prepare heme-depleted media (HDM; see *Materials and Methods*). **b.** Heme levels of indicated media, namely HDM, RPMI with 10% FBS, and HDM with 5µM heme (or Heme repleted media). **c.** Intracellular heme concentrations achieved in cells were incubated in HDM with heme at 0 µM (light grey), 5 µM (dark grey) or 50 µM (black). **d.** Heme concentrations that induced 50% death ($IC_{50}$) in cells expressing GFP alone or K13-GFP. **e.** Effect of indicated media (namely control medium, HDM, HDM + 5 µM heme and HDM + ALA) on the expression of GFP: Western blots and associated bar graphs are shown. **f.** Effect of indicated media (namely control medium, HDM, HDM + 5 µM heme and HDM + ALA) on the expression of KEAP1: Western blots and associated bar graphs are shown. **g.** SDS-PAGE showing differences in trypsin fragmentation profiles between heme-free (lanes 1-4) or heme-bound (lane 5-8) BSA. While trypsin digested the heme-bound BSA into ~45-55-kDa fragments (red arrowheads), much smaller fragments at ~22-, ~34- and ~40-kDa (green arrowheads) were generated by heme-free BSA on trypsin digestion. Lanes 1 and 3 represent heme-free and heme-bound BSA, respectively in the absence of trypsin treatment. Molecular weight standards (M; in kDa) are as indicated. Graphs in panels b-f represent mean from three biological replicates ± SE. Statistical significance calculated by one-way Anova (Tukey's multiple comparisons test). Distinct alphabets represent significant difference at p.adj ≤ 0.05.

### Supplementary Fig. 3

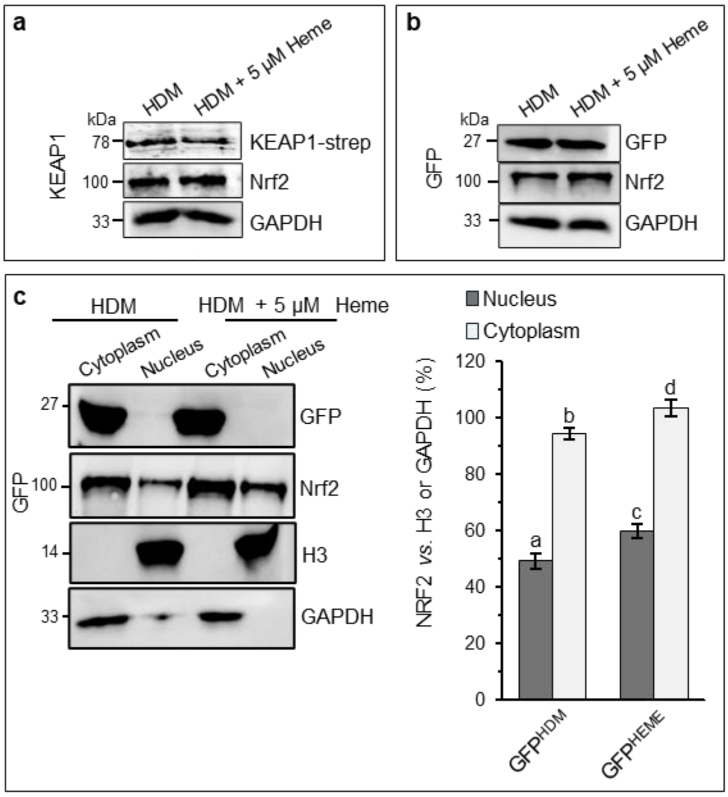


**Supplementary Fig. 3. Effect of heme on the dynamics of Nrf2 in GFP- and KEAP1-expressing Du145 transfectants. a-b.** Western blots showing relative band intensities of either KEAP1-strep (a, top) or GFP (b, top) in transgenic Du145 cells under HDM or 5 µM heme-repleted conditions. Nrf2 (a-b, middle) and GAPDH (loading control, bottom) were also detected by specific antibodies. **c.** Western blots (left) and graphical densitometric quantitation (right) for cytoplasmic and nuclear fractionation of GFP (top blot), Nrf2 (second blot) in $GFP^{HDM}$ cells and $GFP^{HEME}$ cells. Histone H3 (third blot) and GAPDH (bottom blot) validated the purity of nuclear and cytoplasmic fractions, respectively. Molecular weight (in kDa) are as indicated for all blots. Graph represents mean from three biological replicates ± SE. Statistical significance calculated by one-way Anova (Tukey's multiple comparisons test). Distinct alphabets represent significant difference at p.adj ≤ 0.05.

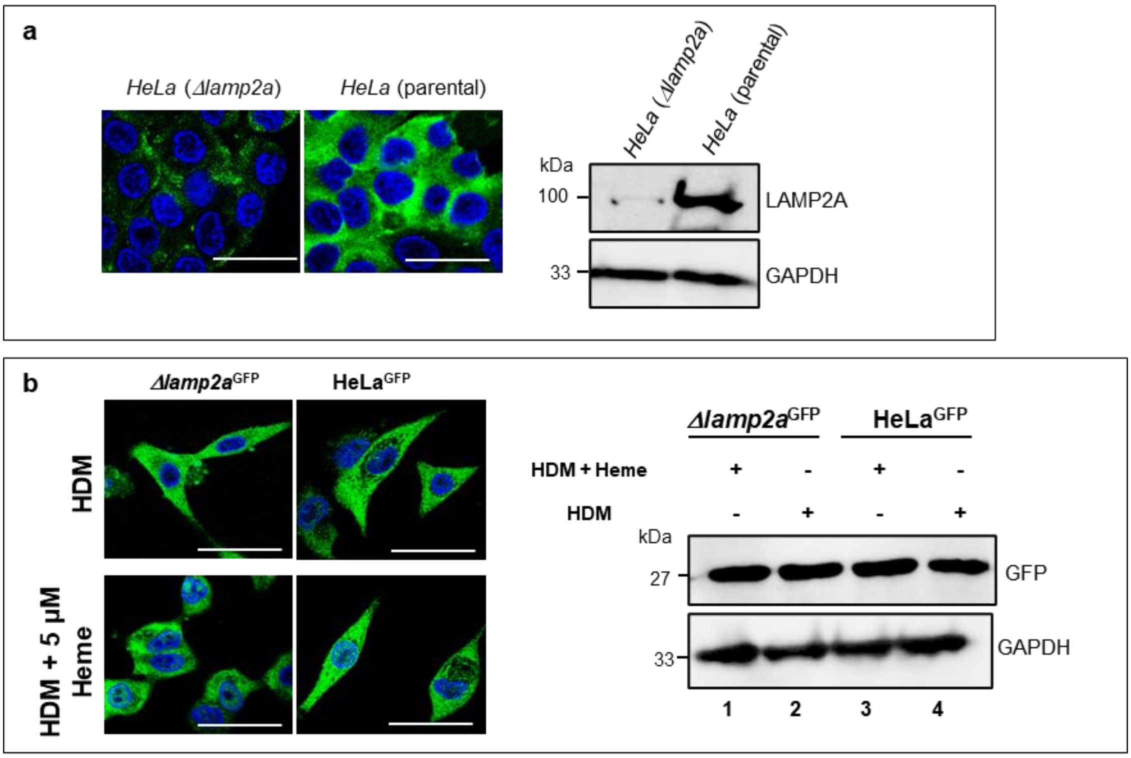


**Supplementary Fig. 4. Characterization of *lamp2a* knockout cells and consequence on GFP expression under HDM or heme-repleted conditions. a.** IFA images (left) and western blot (right) showing no LAMP2A expression in the HeLa *Δlamp2a* cells as compared to the parental HeLa cells. **b.** IFA images (left) showing no change in the fluorescence intensity of GFP between the Δ*lamp2a*$^{GFP}$ cells (left) as compared to the parental HeLa$^{GFP}$ cells (right) under HDM (top) or HDM + 5 μM heme supplemented conditions (bottom). Western blots (right) show no change in the GFP protein levels in transgenic *Δlamp2a*$^{GFP}$ HeLa (lanes 1-2) or parental HeLa$^{GFP}$ cells (lanes 3-4) under HDM (lanes 1 and 3) or 5 μM heme-supplemented (lanes 2 and 4) conditions. Antibodies recognized GAPDH protein as a loading control (bottom blots in a and b). Molecular mass in kDa is as indicated. For all IFA images, nuclei (blue) were stained with Hoechst 33342; scale bar 5 μm.
